# Activation of GPR116/ADGRF5 by its tethered agonist requires key amino acids in extracellular loop 2 of the transmembrane region

**DOI:** 10.1101/2021.04.01.438115

**Authors:** James P. Bridges, Caterina Safina, Bernard Pirard, Kari Brown, Alyssa Filuta, Rochdi Bouhelal, Sejal Patel, Klaus Seuwen, William E. Miller, Marie-Gabrielle Ludwig

## Abstract

The mechanistic details of the tethered agonist mode of activation for adhesion GPCRs has not been completely deciphered. We set out to investigate the physiologic importance of autocatalytic cleavage upstream of the agonistic peptide sequence, an event necessary for NTF displacement and subsequent receptor activation. To examine this hypothesis, we characterized tethered agonist-mediated activation of GPR116 *in vitro* and *in vivo*. A knock-in mouse expressing a non-cleavable GPR116 mutant phenocopies the pulmonary phenotype of GPR116 knock-out mice, demonstrating that tethered agonist-mediated receptor activation is indispensable for function *in vivo*. Using site-directed mutagenesis and species swapping approaches we identified key conserved amino acids for GPR116 activation in the tethered agonist sequence and in extracellular loops 2/3 (ECL2/3). We further highlight residues in transmembrane7 (TM7) that mediate stronger signaling in mouse versus human GPR116 and recapitulate these findings in a model supporting tethered agonist:ECL2 interactions for GPR116 activation.

**Grant support:** This work was supported in part by HL131634 (JPB) from the National Heart, Lung and Blood Institute of the National Institutes of Health.

## Introduction

Adhesion G Protein-Coupled Receptors (aGPCRs) are involved in a variety of pathophysiological processes, from development of the brain and the musculoskeletal system to modulation of metabolism (Olaniru et al., 2019), immune responsiveness and angiogenesis. Important roles in the central and peripheral nervous system have been described (reviewed in Folts et al., 2019), exemplified by the genetic linkage of GPR56/ADGRG1 variants with bilateral frontoparietal polymicrogyria (BFPP) pathology. The capacity of aGPCRs to modulate inflammatory responses has been shown in several contexts (Lin et al., 2017), with leading data on apoptotic cell sensing and phagocytosis (EMR2/ADGRE2, BAI1/ADGRB1), activation of mast cells (EMR2 genetic link to vibratory urticaria) or NK cells (GPR56), and resistance to listeriosis (CD97/ADGRE5). In addition, several lines of evidence suggest key roles for aGPCRs in cancer where they may, for example, relay cellular signaling of mechanical cues (reviewed in Scholz, 2018).

A specific aspect of aGPCR biology is their mechanosensory role, an aspect that may prove to be a general feature of these receptors (Langenhan, 2019). These properties are related to the structure of aGPCRs and their mode of activation that was recently identified. Several methods of activating aGPCRs have been identified, including using endogenous ligands, peptide agonists and small molecules (reviewed in Bassilana et al., 2019). What may be seen as the primary activation mode of aGPCRs is activation via a tethered agonistic peptide (Liebscher et al., 2014; Stoveken et al., 2015). Intracellular autocatalytic cleavage of aGPCRs at the GPS site upstream of the seven transmembrane domain (7TM) generates the NTF and CTF (N-terminal and C-terminal fragments, respectively), with the amino terminal 15 to 27 amino acids of the CTF being referred to as the tethered agonist. The NTF and CTF then re-associate non-covalently during trafficking to the cytoplasmic membrane such that the tethered agonist is buried within the C-terminal portion of the NTF (Araç et al., 2012). Interactions of the NTF with extracellular matrix (ECM) components, or with neighboring cells, serves as an anchor for the receptor. Upon binding of an additional ligand that changes the structural conformation, or generation of tension (e.g. shear stress, stiffness of the ECM), the tethered agonist is released from the NTF thereby activating the 7TM of the receptor. Several studies have shown constitutive basal activity of CTF constructs and activation of full-length aGPCRs with exogenous synthetic peptides (GAP) corresponding to their cognate tethered agonist sequence (Liebscher et al., 2014; Demberg et al., 2015; Müller et al., 2015; Stoveken et al., 2015; Wilde et al., 2016; Brown et al., 2017; reviewed in Bassilana et al., 2019). Some receptors such as CELSR1/ADGRC1, GPR115/ADGRF4, GPR111/ADGRF2 do not possess a consensus GPS cleavage site and are not cleaved. Further, the biological function of GPR114/ADGRG5 and LAT1 (LPHN1/ADGRL1) is not dependent on the cleavage at the GPS site and such receptors may be activated by structural changes not requiring the dissociation of the NTF and the CTF (Langenhan 2019; Scholz et al., 2017; Beliu et al., 2021). In contrast, release of the NTF, concomitant to receptor activation, has been shown for GPR56 upon binding to the ECM laminin 111 and transglutaminase 2 (Luo et al., 2014) and for EMR2 bound to the ECM dermatan sulfate and subjected to mechanical cues (vibration) (Boyden et al., 2016; Le et al., 2019; Naranjo et al., 2020).

However, the molecular details for the activation of aGPCRs by their tethered agonists have not been well described; in part due to difficulty of generating 3D structures of GPCRs. Multiple sequence alignments from previous reports revealed a 25-30% amino acid identity between aGPCRs and class B1 GPCRs (Bjarnadóttir et al., 2007; Stacey et al., 2000). Available 3D structures of class B1 GPCRs (19 structures, 11 X-ray and 8 cryo-electron microscopy (cryo-EM)) in the GPCR DB, www.gpcrdb.org as accessed on January 8^th^ 2020, have provided templates for homology models which could be used to gain insights into the structure of the CTF of aGPCRs. However, it remains unclear as to what extent this knowledge on class B receptors will be directly applicable to the mechanism of aGPCR activation.

For aGPCRs, specific domains of the extracellular region have been crystalized. In a seminal study, the GAIN and Horm domains of two aGPCRs from different subfamilies, LPHN1 and BAI3/ADGRB3 were crystalized (Arac et al., 2012). This study defined the GAIN domain as the important autocatalytic structure, containing the GPS cleavage site, and positioned the tethered agonist within the cleaved NTF (Beliu et al., 2021). Partial NTFs of LPHN3/ADGRL3 and FLRT3 have also been crystalized and, when combined with the FLRT3/UNC5 structure, validated formation of LPHN3/FLRT3/UNC5 trimers in cell interactions (Lu et al., 2015). Crystallization of the full extracellular domain of GPR56 identified new domains in the NTF and suggested hypotheses of how receptor activity is modulated by different NTF variants. CRF1 and GAIN-Horm domain structures were used to model LPHN1 and study its interaction with Teneurin2 (Li et al., 2018). The GLP1R structure was used to model the interaction of CD97 with beta arrestin 1 (Yin et al., 2018). In 2020, the NTF of GPR126/ADGRG6 was crystalized and its structure defined, including the functional implications of two splice variants (Leon et al., 2020). Very recently, Ping and colleagues (Ping et al., 2021) reported the first cryo-EM structures of the glucocorticoid-bound aGPCR GPR97/ADGRG3-G0 complex. Their work provides a structural basis for ligand binding to the CTF of aGPCRs and subsequent G protein coupling, while molecular details of the interaction of the tethered agonist with the CTF remain elusive. Peeters et al. (2016) analyzed activation motifs in GPR112/ADGRG4, comparing conserved functional elements between TMs of class A, class B receptors and this aGPCR. A similar study on GPR56 and ELTD1/ADGRL4 gave further elements on structural transmembrane (TM) motifs that may regulate the activation of most aGPCRs (Arimont et al., 2019). Nazarko et al. (2018) performed a comprehensive mutagenesis screen of LPHN1 to get a more detailed insight into the molecular activation mechanisms at the level of the TM domains. With regard to the tethered agonist peptide, a beta strand conformation has been suggested, followed by a turn element that may be key in receptor activation (detailed in Vizurraga et al., 2020). However, only recently a study explored the modalities of the tethered agonist-mediated activation (Sun et al., 2020). We set out to investigate in-depth the tethered agonist-induced activation of aGPCRs, by NTF removal or exogenous GAP addition, using GPR116/ADGRF5 as an example.

GPR116 is expressed in several tissues and cell types, including alveolar type II epithelial (AT2) cells in the lung. One of its major roles is the regulation of pulmonary surfactant revealed by global knock-out of the gene in mice (Bridges et al., 2013; Yang et al., 2013; Fukuzawa et al., 2013; Niaudet et al., 2015). Cell selective deletion of GPR116 in AT2 cells of adult mice or deletion of the signaling protein GNAQ recapitulated the increased pulmonary surfactant phenotype observed in the global knock-out mice (Brown et al., 2017). Still considered an orphan aGPCR, GPR116 has been proposed to bind surfactant protein D (SFTPD) via its NTF (Fukuzawa et al., 2013). However, evidence supporting a direct role for SFTPD in activating GPR116 is lacking. In order to evaluate the role of the GPS cleavage and the tethered agonist for *in vivo* activation and functionality of GPR116, we generated knock-in mice in which the autocatalytic cleavage at the GPS site is prevented. This type of approach has already been used for other aGPCRs in *C. elegans* and in *in vitro* systems, using mutations immediately upstream or downstream of the cleavage site (Kishore et al., 2016; Prömel et al., 2012; Peeters et al., 2015). Moreover using an approach similar to that developed for PAR receptors (O’Brien et al., 2001), we generated synthetic peptides corresponding to the tethered ligand sequence and mutations within the extracellular loops of GPR116 to identify key residues involved in binding between the tethered ligand and its orthosteric binding site. Together, our findings reveal significant knowledge into the mechanisms underlying the activity of the GPR116 aGPCR *in vivo*.

## Results

### Physiological relevance of the GPS cleavage

GPR116 contains a conserved GPS cleavage site and receptor activation by the corresponding tethered agonist has been demonstrated in previous studies *in vitro*. Here, we set out to investigate the relevance of the GPS cleavage for the physiological functions of GPR116 *in vivo*. With this aim in mind, we first generated a mutant GPR116 H991A construct containing a histidine to alanine mutation at amino acid 991 (Fig1A). Mutation of the histidine residue at this position was chosen as it is conserved in rodent and human GPR116 and is known to abolish autocatalytic cleavage of the GPS in other aGPCRs (Zhu et al., 2019). We expressed wild-type and H991A GPR116 in HEK293 cells and demonstrated by Western blot that the H991A is not cleaved. In particular, while the CTF fragment and full-length proteins were detectable in cells expressing the wild-type construct, only the full-length protein was detectable in cells expressing the H991A construct (Fig1B). The non-cleavable H991A mutant receptor also traffics to the membrane similar to its wild-type counterpart (Fig1B). With respect to signaling, in both calcium transients (Fig1C) and in IP conversion assays (Fig1D), the response of GPR116 H991A to exogenously added agonistic peptide is indistinguishable from that of the wild-type receptor. These results taken together indicate that the H991A mutant is capable of proper trafficking to the membrane, is able to response to exogenous peptide, but is unable to be cleaved and activated by endogenous ligands *in vivo*.

**Figure 1.**
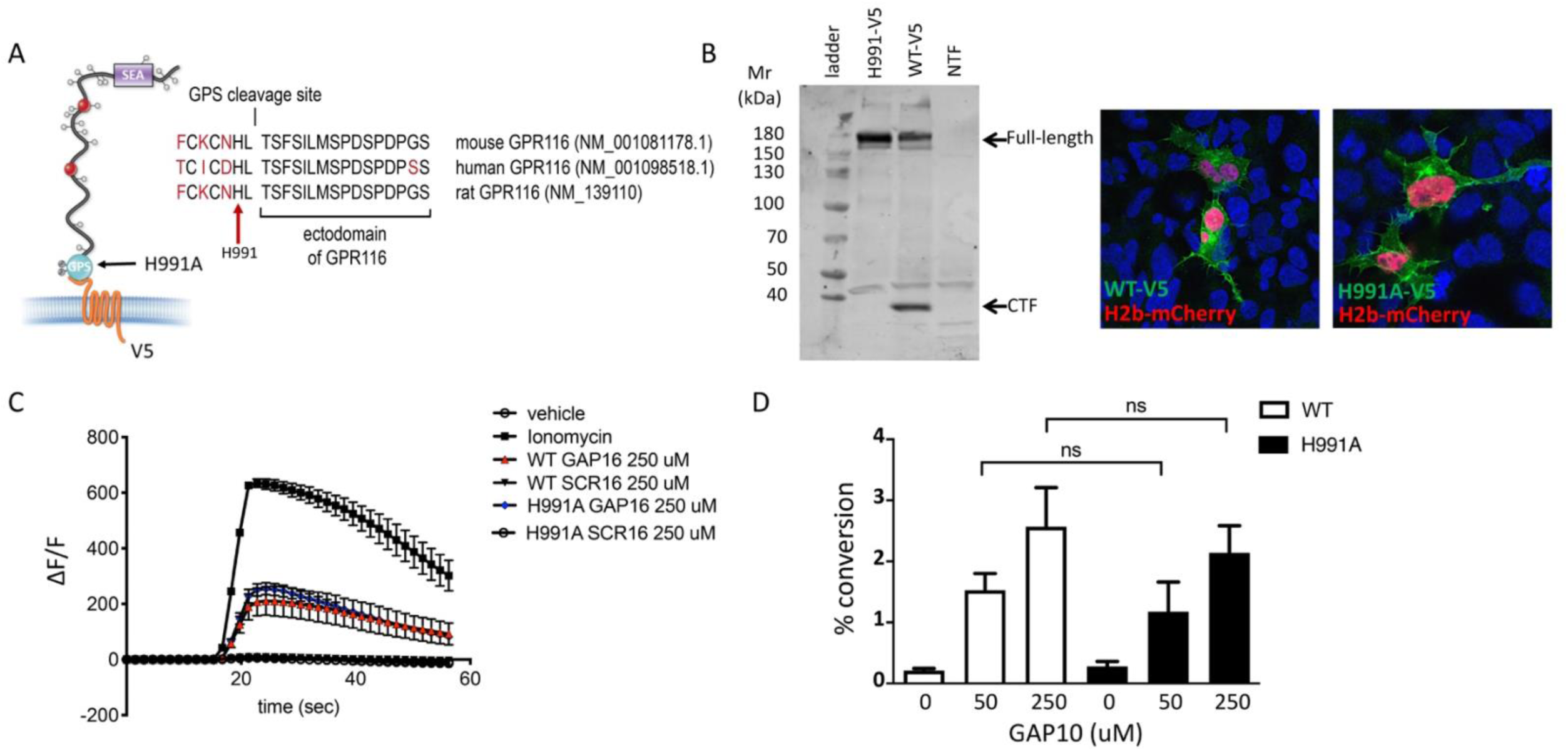
Generation and validation of a non-cleavable GPR116 mutant H991A. **A.** Schematic representation of the H991A mutation in GPR116, details of the sequence at the protein level and gene references. Non-conserved amino acids are highlighted in red. **B.** Transient expression of V5- tagged H991A in HEK293 cells shows no cleavage at the GPS site by Western blot analysis compared to wild-type (WT). Membrane localization of WT and H991A receptors in HEK293 cells by V5 immunocytochemistry. **C,D.** Functional characterization of mouse GPR116 H991A in calcium transient assays (**C**) and in IP accumulation assays (**D**), using GAP16, GAP10 or scrambled peptide (SCR16) as the stimulus (n=3 independent experiments, 3 biological replicates per group). Data are expressed as mean +/- SD (1-way ANOVA for C and D). ns = not significant.

Following functional characterization of the H991A mutation *in vitro*, we introduced the same single amino acid substitution into the GAIN domain of the mouse *Adgrf5* locus via CRISPR/Cas9 gene editing (SupplFig1). Bronchoalveolar lavage fluid (BALF) was obtained from 3 independent lines of 4-week old H991A homozygous mice and compared to BALF obtained from wild-type mice (Fig2A). Interestingly, all three H991A lines contained increased levels (∼65-90umol/kg) of saturated phosphatidylcholine (SatPC) compared to BALF obtained from wild-type mice (∼10umol/kg). Histological analysis of homozygous H991A animals revealed accumulation of eosin-positive material and foamy alveolar macrophages in the distal airspaces. The lungs also demonstrated significant levels of alveolar simplification as noted by increased airspace/tissue density (Fig2B), consistent with the pulmonary phenotype observed in germline *Gpr116*^-/-^ mice (Bridges et al., 2013). We verified functionality of the H991A receptor *in vivo* and found that purified AT2 cells from homozygous H991A knock-in mice responded to exogenous peptide stimulation *ex vivo* (Fig2C,D), generating calcium transient responses similar in magnitude to that previously observed in WT AT2 cells (Brown et al., 2017). Taken together, the H991A mutation prevents cleavage of GPR116 *in vivo* resulting in a pulmonary phenotype similar to that observed in GPR116 knock-out mice. These data demonstrate that while the non-cleavable receptor is fully activated *in vitro* by exogenous peptides corresponding to the tethered agonist sequence, cleavage of the receptor and unmasking of the tethered agonist sequence is critical for GPR116 activation *in vivo*. We next investigated the mechanisms of tethered agonist-mediated GPR116 activation in detail.

**Figure 2.**
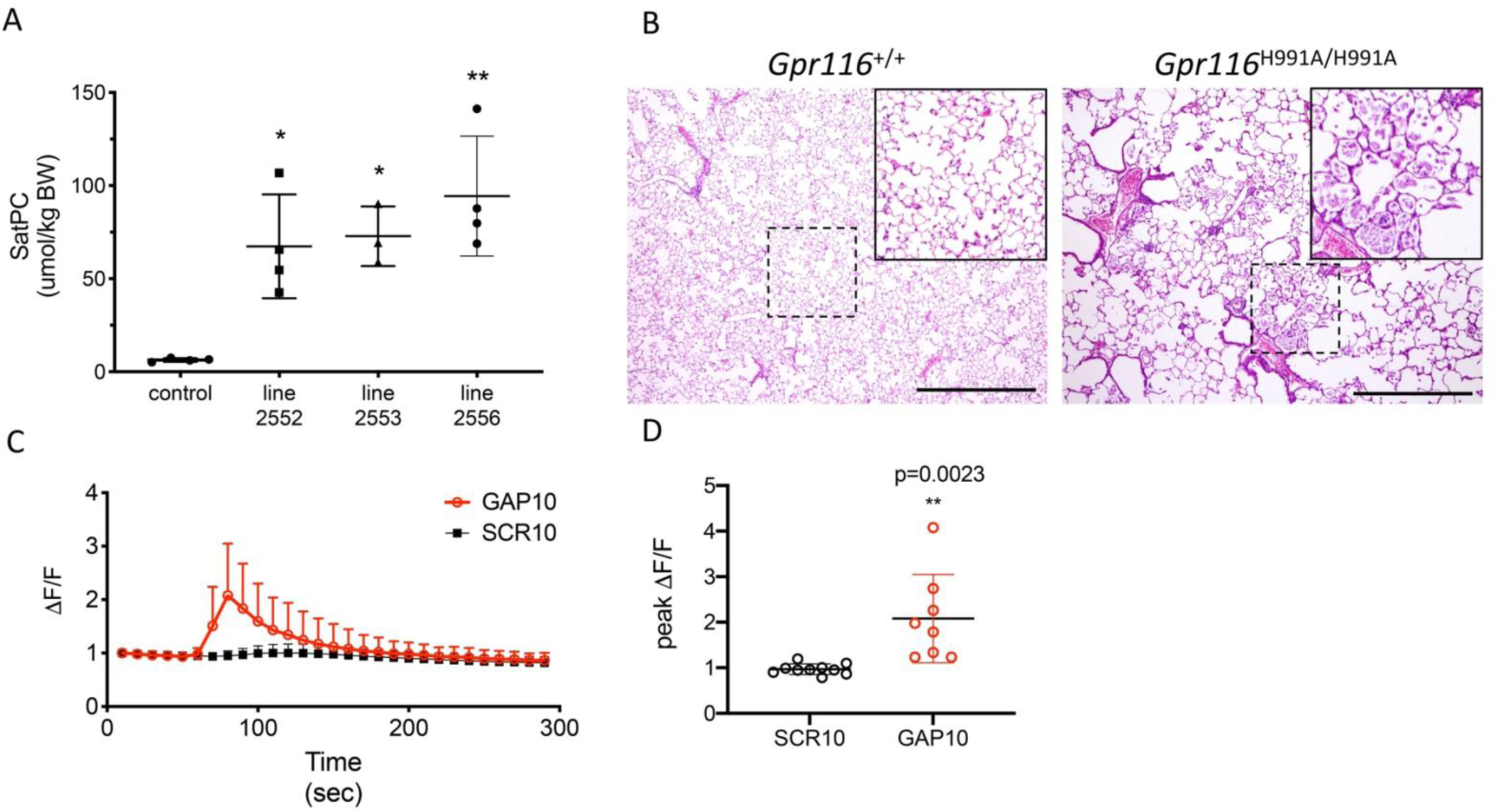
Cleavage at the GPS site is required for GPR116 function in vivo. **A.** Content of saturated phosphatidylcholine (SatPC) in the bronchoalveolar lavage fluid (BALF) of 4 week old wild-type control and H991A homozygous knock-in mice (n=3-4 mice per group). Data are expressed as mean +/- SD (1-way ANOVA). ** P<0.005, * P<0.05. **B.** Representative histology of 4.5 month old wild-type and homozygous H991A knock-in mice. Note accumulation of pulmonary surfactant (inset) and alveolar simplificaiton in H991A knock-in mice compared to wild-type control. **C**. GAP10-induced calcium transients in AT2 cells of H991A homozygous knock-in mice (n=3 independent experiments, 3 biological replicates per group). **D.** Peak calcium responses in H991A homozygous H991A AT2 cells treated with SCR10 or GAP10 (n=3 independent experiments, 2-3 biological replicates per group).

### Identification of key amino acids in the tethered agonist required for GPR116 activation

The functional role of the tethered agonist in GPR116 has been shown previously *in vitro* using the C-terminal fragment (CTF) construct which exhibits strong basal activity due to the presence of unmasked tethered agonist at the amino terminus (Brown et al., 2017). In our previous study, mutation of the tethered agonist amino acids to alanines using sequential steps of 3 amino acids for each mutant highlighted the role of this N-terminal sequence in receptor activation. The minimal exogenous peptide capable of activating GPR116 was defined as GAP9, corresponding to the 9 most N-terminal amino acids of the tethered agonist. To identify the key amino acids critical for activation, we mutated the 12 most N-terminal amino acids individually to alanine using mouse GPR116 CTF (mCTF) as the parent construct (Fig3A). All mutants showed comparable expression levels to that of mCTF (Fig3B). To evaluate the basal activity of the different CTF constructs we used the IP1 conversion assay. We found that 3 mutants showed complete inhibition of IP1 formation: F995A, L998A, M999A (Fig3C). Significant effects were also observed for I997A, D1002, S1003 and P1004 mutations which showed diminished, but not completely abolished, activity. We then determined whether the inactive mutants would function in a dominant negative fashion or if they could still be activated by exogenous GAP peptides. To this end, we first demonstrated that the wild-type reference mCTF construct could be activated above basal levels (super-activated) with exogenous GAP peptide, and that inclusion of an N-terminal FLAG tag did not affect the response (Suppl Fig2A, B). Next, we evaluated the different mCTF single alanine mutants for their ability to be activated by the exogenous ligand. In all cases GAP14 strongly activated the mutant receptors, with IP1 levels accumulating to levels equivalent to or exceeding those observed with the reference wild-type construct (Fig3D). Interestingly, for alanine mutant constructs that lost their basal activity, such as F995A, GAP14 could completely rescue the activity of the mutated tethered agonists. Constructs in which the basal activity was not affected by the mutations were also super-activated by GAP14 similar to the reference mCTF construct. These data are consistent with our previous results with sequential 3 amino acid mutants (Brown et al., 2017) and further show that the tethered agonist single point mutants are still capable of activation by GAP14, demonstrating that the ligand binding site remains available and accessible in these mutants. Furthermore, the data indicate that F995, L998, M999 are the key amino acids within the tethered agonist sequence involved in receptor activation.

**Figure 3.**
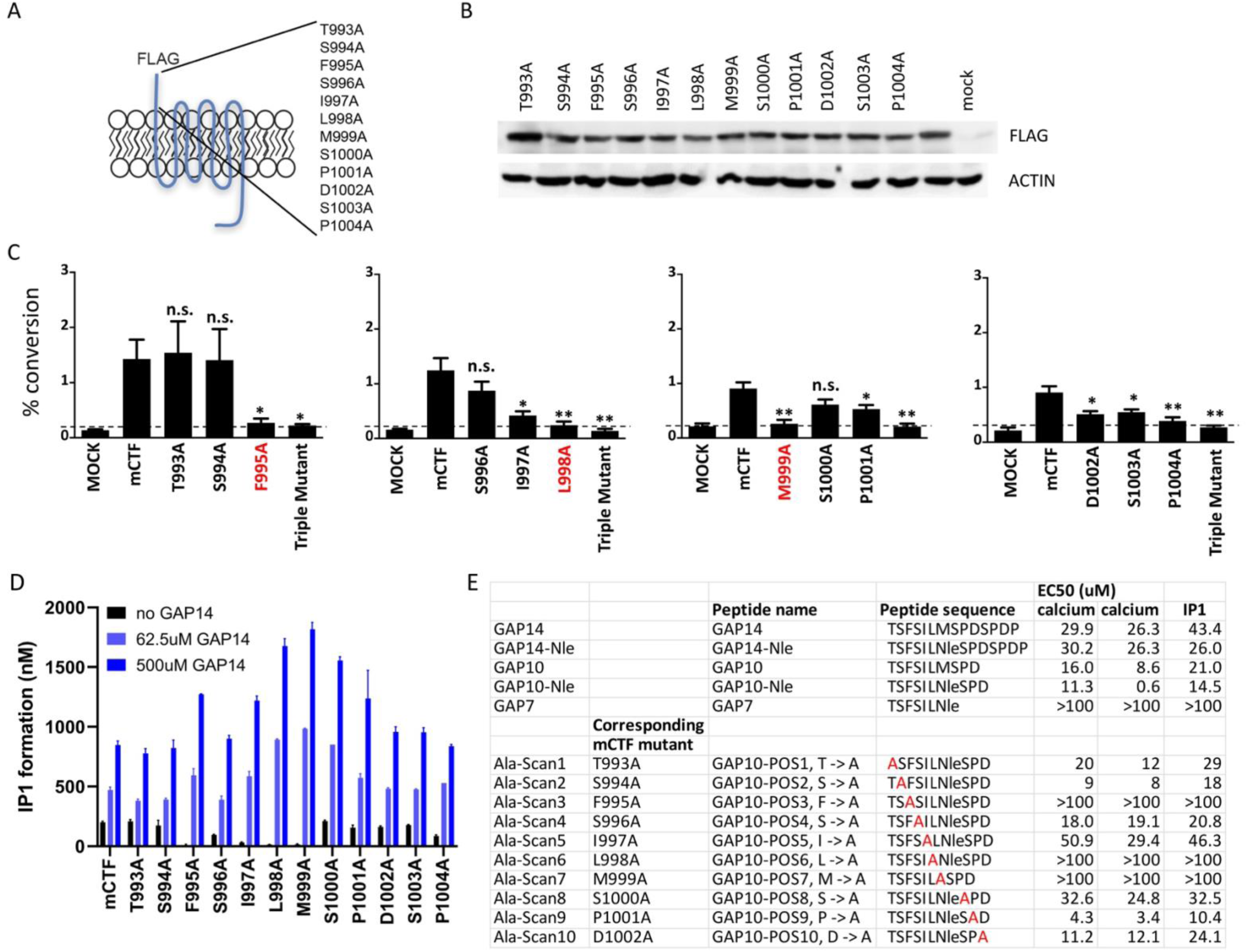
Identification of amino acids in the tethered agonist essential for GPR116 activity. **A.** Design of the 12 mGPR116 CTF (mCTF) ECD mutants. Alanine scan of individual residues in the N-terminal sequence. **B.** Expression of the mCTF ECD mutants transiently expressed in HEK293 cells. Constructs were detected by Western blot of whole cell lysates using an anti-FLAG antibody. **C.** Signaling of the mGPR116 CTF ECD mutants. Constructs of the ECD alanine scan and corresponding triple mutants were transiently expressed in HEK293 cells and basal activity was measured as % IP1 conversion (n=4-5 independent experiments, 3 biological replicates per group). Data are expressed as mean +/- SD (1-way ANOVA). ** P<0.005, * P<0.05. The 3 single point mutants with the strongest effects are highlighted in red. **D.** Exogenous GAP14 treatment rescues mCTF ECD mutants with inactive tethered agonist and super-activates constructs exhibiting basal activity. Constructs were transiently expressed in HEK293 cells and stimulated with GAP14. IP1 accumulation was measured to evaluate basal and GAP14-induced activity of the receptor. Basal activity of mCTF (dashed line) was used as the reference. **E.** Activation of full-length mGPR116 in stable expressing cells (HEK293 clone 3C) with exogenous GAP10 peptides that were sequentially mutated to alanine at each position. Receptor activation was measured via calcium transient assays (n=2 independent experiments) and IP1 accumulation assays (n=1 experiment).

We next set out to confirm the role of critical amino acids in the tethered ligand using mutated versions of exogenous agonist peptides in the context of the full-length receptor. We used GAP10 as the template for these studies, as it is more active on the murine receptor than GAP14. In order to rule out potential artifacts resulting from oxidation, the methionine corresponding to position 999 in the mCTF was replaced by norleucine (Nle). The activity of GAP10 Nle was indistiguishable from that of parental GAP10 (Fig3E). IP1 accumulation and calcium flux assays were carried out to measure the activity of GAP10 variants (Fig3E). GAP10 peptides corresponding to the mutated CTF constructs F995A, L998A and M999A failed to activate the full-length receptor, confirming the results shown above using the mCTF constructs with basal activity. Similarly, peptides corresponding to the other singly mutated mCTF constructs showed activities comparable or weaker to parental GAP10 in agreement with the results obtained with the mCTF constructs. Importantly, we confirmed that the mutant GAP10 peptides F995A, L998A, M999A also failed to activate the human full-length receptor, whereas parental GAP10 was active (Suppl Fig3). These results not only identify the critical residues of the tethered agonist sequence, but also show that the functionality of the 3 most critical amino acids in the GPR116 tethered ligand is conserved across species.

### Identification of key amino acids in the 7TM domain of GPR116 involved in tethered agonist-mediated receptor activation

With strong evidence of the residues in the tethered ligand required for receptor activation and the confirmation that GPR116 utilizes the tethered agonist signaling mode, we set out to determine the region of the receptor to which the tethered agonist binds to facilitate receptor activation. Importantly, CTF constructs with a FLAG or mCherry tag attached to the N-terminus retain full basal activity. This suggests that the tethered ligand does not need to penetrate deeply into the transmembrane domains of the receptor for activation. Rather, this suggests that the critical contact sites for the tethered ligand are located on the outside facing surface of the protein, within or proximal to the extracellular loops, similar to the mode of activation of PAR1 (Seeley et al., 2003).

We therefore performed a comprehensive alanine scan of 51 amino acids located within the three predicted ECLs, including a few amino acids extending into the TMs (Fig4A). Three native alanines located in this selection were mutated to valines. For this scan, we used a mouse GPR116 CTF construct that is deficient in basal activity yet responsive to GAP-mediated activation as the parent construct. This construct, initially described in Brown et al. (2017), has the three N-terminal amino acids of the tethered agonist sequence (Thr993, Ser994, F995) mutated to alanines and is refered to hereafter as mCTF ‘basal activity deficient’ mutant, or mCTFbax. Data from this alanine scan identified 5 constructs that were unresponsive to GAP14- mediated activation, yet exhibited expression levels comparable to the parent construct: Y1158A, R1160A, W1165A, L1166A and T1240A. (Fig4B,C). The snake plot in Fig4A depicts these amino acids as pink circles. Four of these key amino acids are located in ECL2 (Y1158, R1160, W1165 and L1166A) while T1240 is located at the top of TM6, adjacent to ECL3. Of note, the disulfide bridge between the top of TM3 and ECL2 in GPCRs is known to be required for proper receptor conformation. We verified that alanine mutation of C1088, which is predicted to interact with C1164, indeed eliminated receptor activation by the agonist peptide (Suppl Fig4). All other mutants tested showed comparable activation to the parent construct or partial activation (Suppl Fig4 and data not shown).

**Figure 4.**
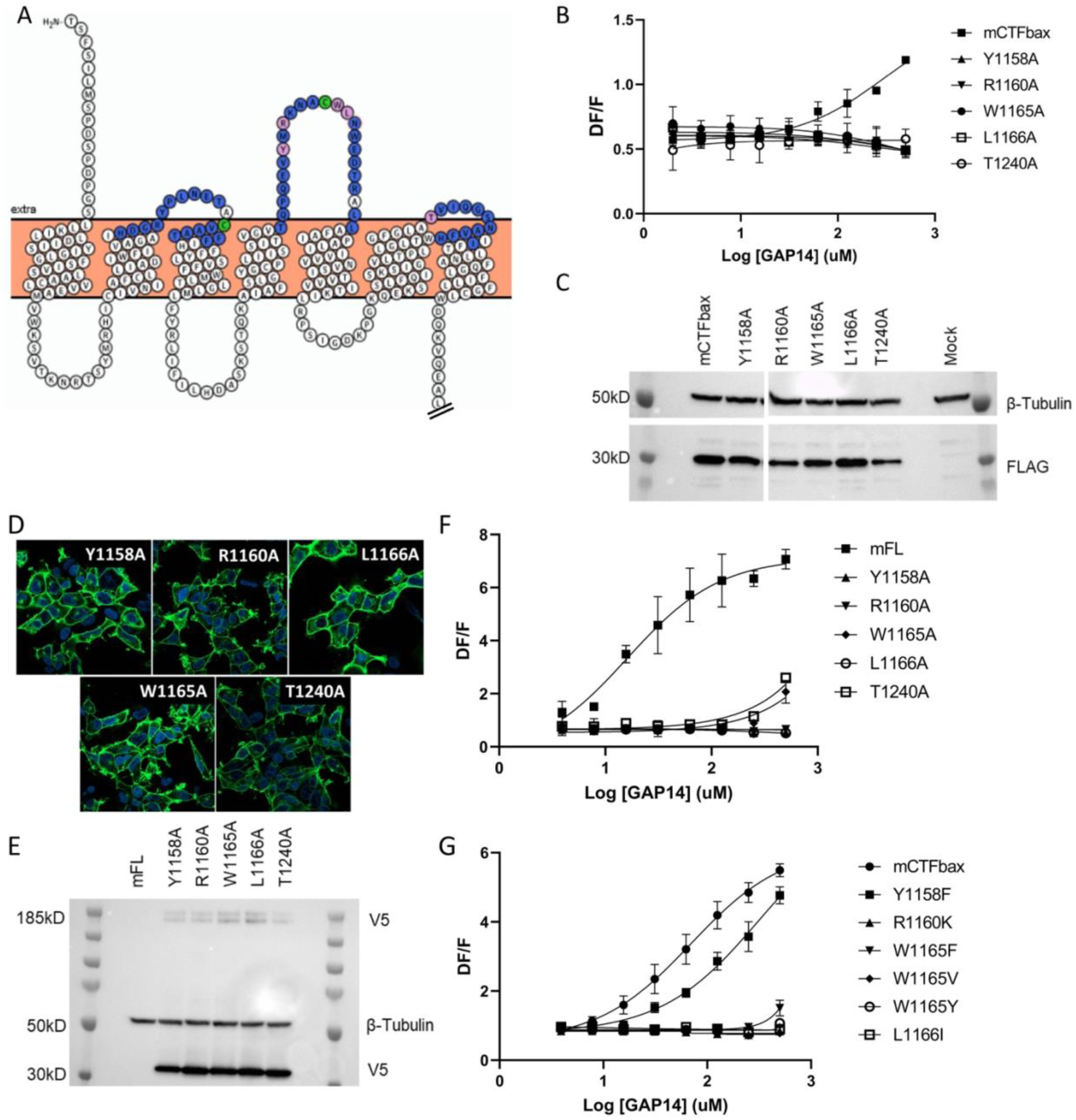
Identification of key ECL amino acids involved in GPR116 activation by the tethered agonist. **A.** Snake plot model of mouse GPR116 CTF. ECL residues individually mutated in the alanine scan of mouse GPR116 CTF constructs tested for functional activity are colored in blue or pink; the five residues shown in pink were identified as strongly modulating receptor activity. The two cysteine residues forming the disulfide bridge are shown in green. The Ct tail was truncated for visualization purposes as indicated by double hash lines. **B.** Alanine scan of ECLs in mCTFbax, a CTF construct without basal activity. Calcium transient assays were performed in transiently transfected HEK293 cells with exogenous GAP14 as the stimulus. mCTFbax mutants that were completely inactive following GAP14 stimulation are shown. Data are from 3 experiments, each performed in quadruplicates. See Suppl Fig4 for further alanine scan data, including mutants with no effect. **C.** Expression of the mCTFbax mutants not activated by GAP14, in transiently transfected HEK293 cells. Proteins were detected by Western blot using an anti-FLAG antibody. **D.** Membrane expression of the mGPR116 full-length (mFL) ECL mutants assessed by V5 tag immunocytochemistry upon stable expression in HEK293. **E.** Expression of mFL ECL mutants stably expressed in HEK293 cells. Constructs were detected by Western blot using an anti-V5 antibody; the major band (∼30kD) corresponds to the cleaved CTF and the minor bands at high molecular weight (∼185kDa) correspond to the full-length receptor. The mFL WT stably transfected clone 3C is not V5 tagged. **F.** GAP14-induced calcium transients of mFL mutants for 5 key ECL residues, stably expressed HEK293 cells, as compared to the WT mGPR116 3C clone. Data shown are from two independent experiments performed with biological quadruplicates. **G.** Mutation of key ECL residues in the mCTFbax construct to amino acids with functional relevance. mCTFbax and mutants thereof were stably expressed in HEK293 cells and GAP14-induced calcium transients were measured. Data are from 3 experiments each performed in quadruplicate.

The importance of the 5 critical ECL residues identified was further validated in corresponding mGPR116 full-length (mFL) constructs, in mCTF constructs with basal activity, as well as in hGPR116 full-length (hFL). Importantly, these residues are conserved between human and mouse (Suppl Fig5). All 5 point mutants of mFL showed membrane localization (Fig4D) and expression levels similar to that of the wild-type receptor (Fig4E) but could not be activated by exogenous GAP14 (Fig4F). Similarly, these 5 alanine mutations introduced into hFL abolished the response to GAP14, while not affecting protein or plasma membrane localization (Suppl Fig6). Replacing these key residues by alanines in a mCTF construct with high basal activity also strongly decreased or completely abrogated the constitutive activity of the mCTF, without affecting expression levels (Suppl Fig7).

In a related series of experiments we aimed at elucidating whether conservative amino acid replacements within these key residues in mCTFbax would restore receptor activation of the single point alanine mutants. Introducing phenylalanine instead of alanine for Y1158 indeed allowed for receptor activation, albeit with lower potency for the GAP14 agonist peptide (Fig4G). Conservative exchanges for the three other critical amino acids still yielded inactive receptors (R1160K, W1165F, W1165V, W1165Y, and L1166I) all yielded inactive receptors. These mutated receptors showed protein expression levels comparable to the parental mCTFbax construct, with the exception of W1165F and W1165Y, which showed a partial reduction in expression (Suppl Fig8). Our data confirm a predominant role of ECL2 and highlight a specific residue at the top of TM6 / ECL3.

### Species-specific features involved in GPR116 responsiveness to the tethered agonist

The amino acids identified in the CTF that are critical for receptor activation are all conserved between murine and human GPR116 (Fig 4, Suppl Figs5-8). Yet, GAP peptides elicit a more potent response on the mouse compared to the human receptor (Brown et al., 2017), suggesting that additional residues are responsible for this difference in potency. We set out to identify the amino acids relevant for this differential responsiveness to provide additional important insight into the mechanisms underlying GPR116 activation..

We first generated a human GPR116 CTF construct in which the 6 N-terminal amino acids of the tethered agonist were deleted. This construct, referred to as hGPR116 CTFbax (hCTFbax), has no basal activity, analogous to the mCTFbax construct (Fig5A, inset). As expected, the hCTFbax construct responded to GAP14 activation, in IP1 (Fig5A) and calcium flux assays (Fig5B). Similar to observations with the full-length receptors, the hCTFbax is less responsive to GAP14 than the mCTFbax receptor (Fig5B). These constructs were used in subsequent experiments to pinpoint key amino acid residues relevant for this differential responsiveness. With this aim we exchanged specific non-conserved amino acids in the human receptor to the corresponding murine residues, focusing on amino acids at the surface of the receptor in the N-terminus and around ECLs (Fig5C). We mutated small clusters of amino acids as summarized in Fig5D, and also introduced the single mutation G1011K, representing a non-conservative amino acid difference between human and mouse sequences in proximity to the extracellular surface. All mutants expressed well (Suppl Fig9) and were activated by GAP14. Surprisingly, only the G1011K mutation changed the response characteristics of the resulting receptor so that it resembled that of mCTFbax (Fig5E). Position 1011 is located at the top of TM1 and immediately downstream of the tethered agonist.

**Figure 5.**
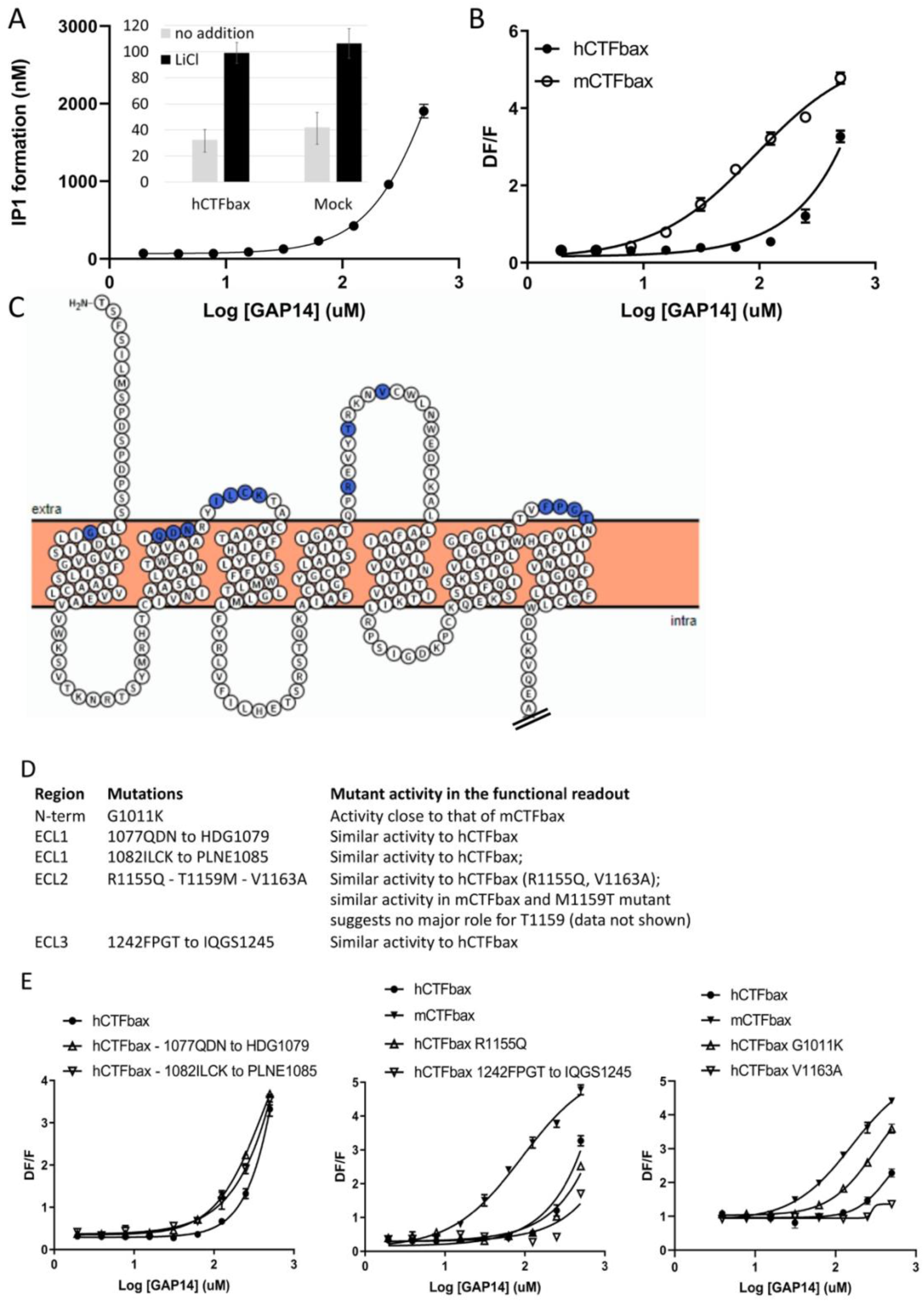
Evaluation of the role of ECL amino acids not conserved between human and mouse GPR116. **A.** Characterization of hCTFbax, a CTF construct without basal activity and that can be activated by GAP14. The hCTF with deletion of the N-terminal 6 amino acids was stably expressed in HEK293 cells and stimulated with GAP14 in an IP1 accumulation assay (n=3 independent experiments with duplicates). The basal level of IP1 formation in presence of LiCl is similar to that of parental HEK cells (inset). **B.** Stable HEK293 populations for mCTFbax and hCTFbax were stimulated with GAP14 and analysed for induction of calcium transients. Data are from n=6-9 repeat experiments with duplicates. **C.** Overview of the mouse-specific residues introduced in the hCTFbax construct. Snake plot of the human GPR116 CTF (including the full tethered agonist sequence); amino acids highlighted in blue, located towards to extracellular side, are not conserved between human and mouse GPR116. The corresponding mouse sequence was introduced in human GPR116. The Ct tail was truncated for visualization purposes as indicated by double hash lines. **D.** Details of the residues exchanged in the hCTFbax with the corresponding mouse sequence. Each cluster of changes was tested for its signaling capacity and the functional assay outcome is described. **E.** GAP14-induced calcium transients were not further increased in the mutants as compared to the reference construct hGPR116 CTFbax with the exception of G1011K (bottom panel). Constructs were stably expressed in HEK293 cells and tested in duplicates. Data were confirmed at least in one repeat experiment (data not shown).

In parallel, while analyzing the potential binding mode of a low molecular weight antagonist specific for the murine receptor discovered in house at Novartis we began considering additional other amino acids deeper in the transmembrane domains of the receptor that might be responsible for the species differences in efficacy. These studies identified a potential binding site containing non-conserved amino acids between mouse and human. We hypothesized that these amino acids might mediate the strong mouse-specific aspects of receptor activation as well as inhibition by the antagonist. We identified 10 mouse-specific residues that may be involved in this putative murine-specific binding site (mBS) (Fig6A) and introduced all 10 residues into human GPR116, generating a hCTFbax mBS chimera (Suppl Fig10A). This construct exhibited an enhanced responsiveness to GAP14, similar to that of mCTFbax (Fig6B). We then mutated individually each amino acid of the mBS. Interestingly, introducing the A1254T or V1258A mutations into the hCTFbax construct resulted in the increased activity of GAP14 (Fig6C), similar to the hCTFbax mBS and G1011K constructs. Other non-conserved amino acids in the N-terminus and ECLs appeared to be irrelevant with respect to activation. Expression of these mutants was verified by V5 immunocytochemistry (Suppl Fig10B). Taken together, our data suggest that K1011, T1254 and A1258 in mouse GPR116 are responsible for the increased potency of GAP14 towards the mouse receptor as compared to human GPR116.

**Figure 6.**
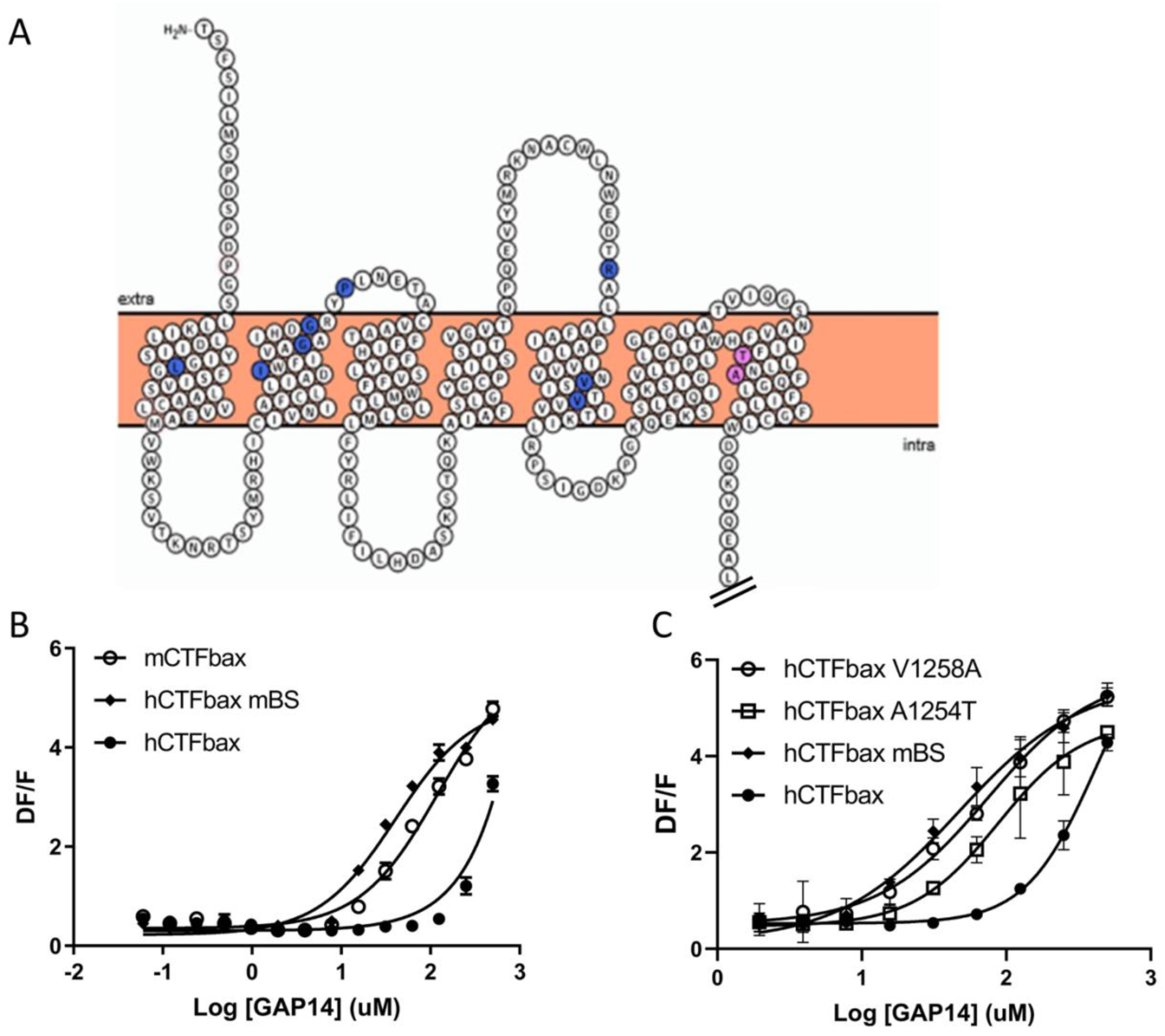
Engineering the predicted mouse GPR116 binding site (mBS) into human GPR116 leads to increased peptide activation. **A**. Snake plot of the mouse GPR116 CTF highlighting mouse-specific amino acids in the putative binding site in blue and pink. Amino acids responsible for mGPR116-specific aspects of receptor activation are shown in pink. The Ct tail was truncated for visualization purposes as indicated by double hash lines. **B.** Stable HEK293 populations for m/hCTFbax and hCTFbax mBS were stimulated with GAP14 and analysed for the induction of calcium transients. A representative graph of 9 repeat experiments is shown. **C.** Characterization of the key amino acids transmitting the effects of the mouse binding site sequence. hCTFbax single mutants were compared to hCTFbax and hCTFbax mBS in calcium transient assays. Stable HEK293 populations were stimulated with GAP14 (n=3 independent experiments each performed in duplicate).

### Identification of a mouse-specific activating peptide with enhanced activity

While studying the mechanisms involved in tethered ligand activation of GPR116, we discovered a GAP10 variant with a threonine to proline substitution in the first position (PSFSILMSPD), named GAP10-Pro1, that was more potent in activating mFL and mCTFbax than GAP10 and GAP14, with little to no effect on hGPR116 (FL or CTFbax). Thus GAP10-Pro1 acts as a highly potent, mouse-specific agonist peptide (Fig7A, left panels versus right panels). Next, we asked whether the same mouse amino acids that confer increased responsiveness to exogenous GAP14 activation in the human receptor would also enable response to GAP10-Pro1. Indeed, the A1254T and especially V1258A hCTFbax constructs showed a significant response to GAP10-Pro1, albeit not fully equivalent to hCTFbax mBS (Fig7B, bottom panel). This study with a modified peptide confirms that we have identified 2 important amino acids involved in specific aspects of mouse GPR116 activation as compared to the human receptor. In contrast, the G1011K mutant did not robustly respond to GAP10-Pro1, similar to parental hCTFbax (Fig7C). This latter obervation suggests that the proline in the novel peptide interacts with residues distinct from amino acid 1011. Moreover, in addition to increasing our mechanistic understanding of the mode of tethered agonist function and species differences between human and murine receptors, this newly discovered agonist will provide a novel reagent to explore additional aspects of the *in vivo* function of GPR116.

**Figure 7.**
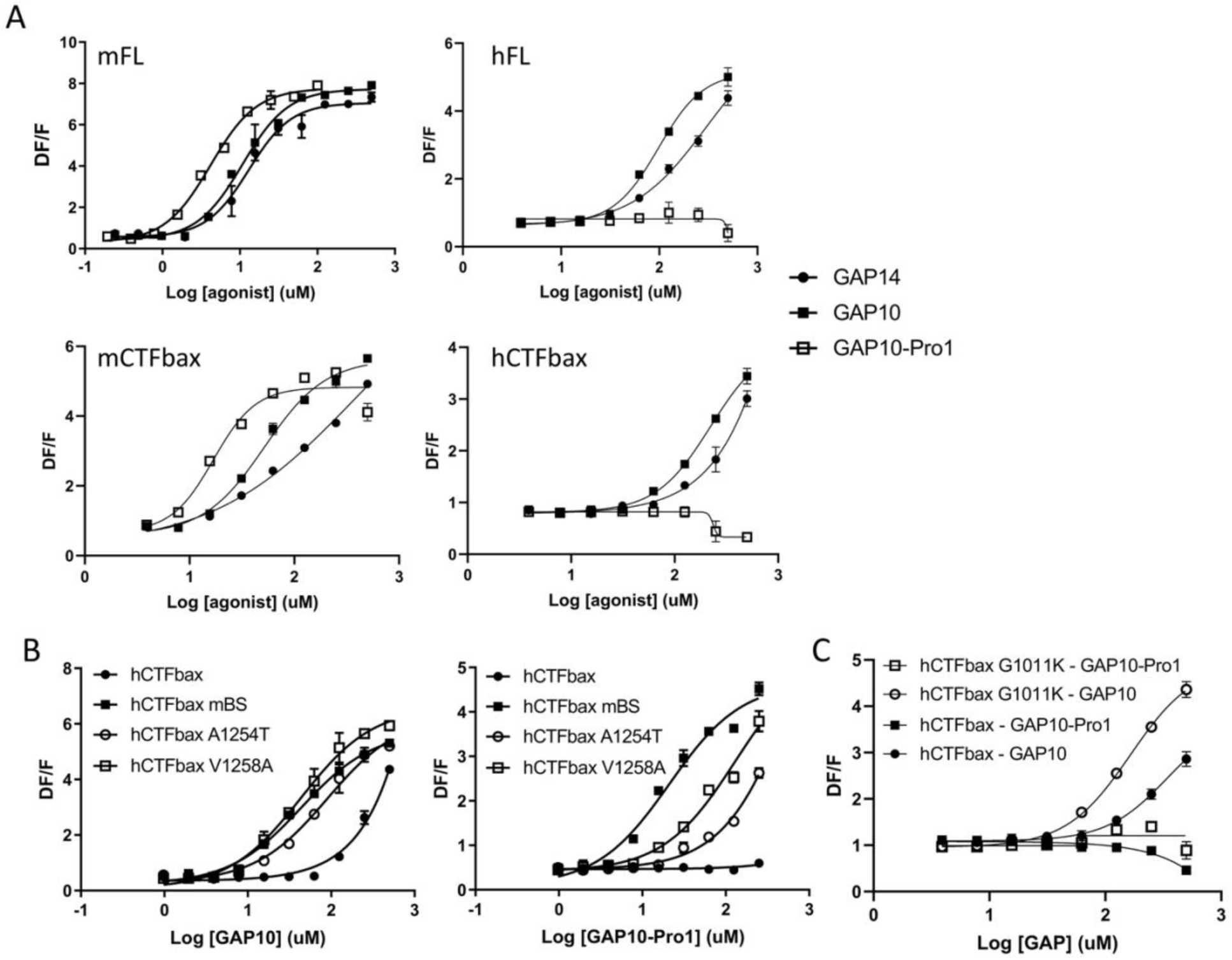
Characterization of a mouse-specific agonistic peptide. **A.** GAP10-Pro1 only activates mouse GPR116. Full-length mouse and human GPR116 (mFL clone 3C and hFL clone A6, respectively), mCTFbax and hCTFbax, all stably expressed in HEK293 cells, were stimulated with GAP14, GAP10 and GAP10-Pro1. Calcium transients are measured, in biological duplicates, as a signaling readout. **B.** Identification of the key amino acids mediating the mouse-specific effects in the mouse binding site. The hCTFbax construct, the hCTFbax mBS chimera and the single point mutants A1254T and V1258A in hCTFbax were stimulated with GAP10 or GAP10-Pro1 upon stable expression in HEK293 cells. Their activation was measured in a calcium transient assays, in biological duplicates. Data are representative of at least two independent experiments with identical results. **C.** Calcium transients evoked in HEK293 cells stably expressing hCTFbax or the mutated construct G1011K upon stimulation with GAP10 or GAP10-Pro1. Data are representative of three independent experiments with identical results.

### Modeling of human GPR116

A receptor model for the CTF of human GPR116 was initially developed using the crystal structure of the human CRF1 receptor as a template. We used the data described in the present report to refine the model. Notably, the five key amino acids in the ECLs that form suggested contact points for the tethered ligand were used to provide constraints and to include the ECLs in the modeling. In addition, the key roles of positions 1254 and 1258 in conferring increased signaling by defining an antagonist binding site for the murine receptor allowed further refinement of the model. The current working model includes the predicted primary binding site comprising these two key amino acids responsible for the increased potency of GAP14 observed in the murine receptor (Fig8A). Fig8B shows a detailed view of ECL2 with key amino acids that mediate receptor activation. A zoom around ECL3 highlights T1240 at the top of TM6, demonstrated in our studies to be important for signaling (Fig8C).

**Figure 8.**
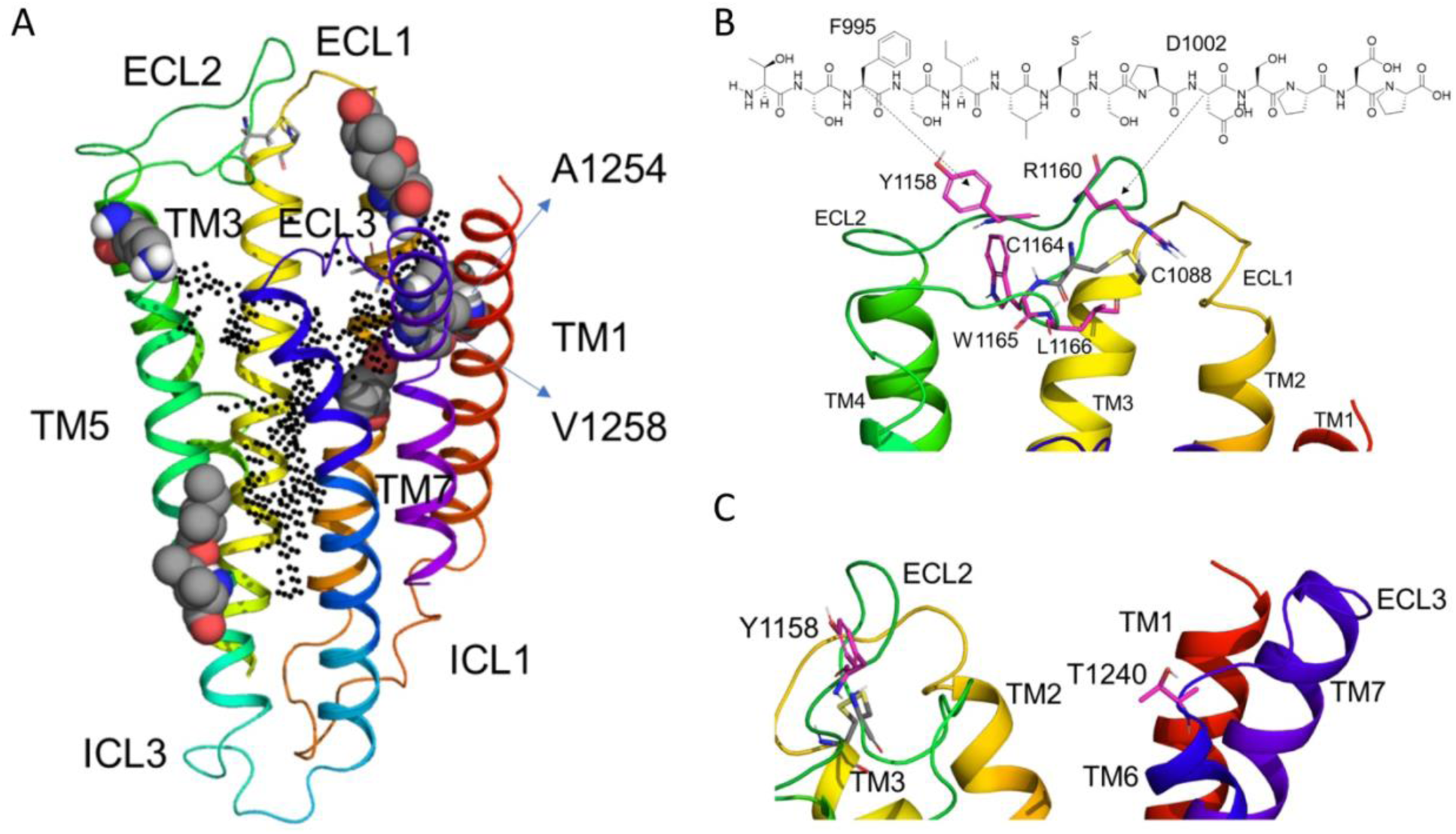
hGPR116 7TM homology model. **A.** Homology model of human GPR116 7TM. The cloud of black dots defines the putative main pocket of the hGPR116 CTF as identified by SiteMap and corresponding to the CRF1R agonist binding site. The mouse-specific amino acids included in the potential antagonist binding site are displayed as spheres and include A1254 and V1258, which generate a stronger activation response upon mutation to the mouse sequence. The residue G1011 involved in stronger mouse GPR116 activation is at the top of TM1 (not shown). Amino acid numbering with respect to the mouse CTF is shown. **B**. Focus on ECL2 highlighting putative key amino acids for GPR116 activation and possible interactions with the tethered agonist drawn on top of the 7TM structure. **C**. Zoom around opposite side of ECLs 2 and 3 highlighting an amino acid at the top of TM6, T1240, important for receptor activation.

The model was further used to illustrate the mode of activation of GPR116 mediated by the tethered peptide interacting with ECL2. We analysed *in silico* how these ECL2 amino acids might interact with the 7TM domain by sampling their different rotamers and selecting the most abundant ones, followed by a local energy minimization to relieve steric clashes within the protein. The activity of the Y1158F mutant in a functional assay had suggested a role for the phenol ring of the tyrosine (Fig4G). This side chain has a high probability of interactint with the phenylalanine in the tethered agonist sequence (F995). It also appears that receptor activation is very sensitive to the nature of the basic side chain at position 1160 as mutation of R1160 to lysine results in lack of response to GAP14. Interestingly, one of the rotamers of the arginine could form salt bridges with both E1169 and with the tethered agonist, presumably D1002. According to this model the change from the arginine with a Y shape to a lysine with a rod-like shape disrupts these suggested interactions and leads to the loss of activity. This is consistent with the observation that the minimal activity of GAP9 is boosted by addition of the aspartate in GAP10, corresponding to D1002 (Brown et al., 2017).

Replacement of W1165 with a phenylalanine, valine or tyrosine significantly reduced the activation potential of the receptor. This suggests a role for the -NH group and that the size of the aromatic ring may be relevant. The most probable interactions of W1165 are with the ECL2 and the TMs rather than with the tethered ligand. Of interest, on top of Van der Waals interactions, the following possible hydrogen bonds were noted: a hydrogen bond with the backbone of V1089 (top of TM3) or T1152 (top of TM4) or with the -OH group in the side chain of T1152. The latter represents an interesting possibility, as it would be specific for the ADGRFs. An interaction with the side chain of H1096 (TM3) would be a conserved mechanism in all aGPCRs, but mutation of the histidine in top of TM3 has no effect in LPHN1 (Nazarko et al., 2018). Surprisingly, replacing L1166 with an isoleucine, as is observed in some aGPCRs, was not tolerated in GPR116, suggesting a specific role of this amino acid. Potential interaction partners for this residues are not defined. This analysis suggested possible tethered peptide:ECL interactions which we integrated into the receptor visualization. In Fig8B, the tethered agonist is represented as an exogenous peptide interacting with ECL2 for activation of GPR116. This scheme summarizes key observations from our study and represents the most likely mode of tethered peptide:ECL interaction required for receptor activation. Moreover, our studies combined with the molecular modeling defines a novel mechanism for GPCR activation.

## Discussion

In this study, we demonstrate that cleavage of GPR116 at the GPS is required for activation and function of the receptor *in vivo*, but dispensable for *in vitro* activation with an exogenous agonistic peptide. Through a detailed mutational analysis of both human and mouse GPR116 and activating peptides we further identified key amino acids in the tethered agonist sequence, in the extracellular loops and the 7TMs of GPR116 that underlie the mechanism of GPR116 signaling.

While great strides have been made in delineating the mechanisms of activation of aGPCRs *in vitro*, the mode of activation of adhesion GPCRs *in vivo*, including GPR116, is not well understood. Through generation and characterization of a cleavage deficient mutant, H991A, we show that cleavage at the GPS site is required for GPR116 activity and modulation of pulmonary surfactant levels *in vivo*, which validates a tethered agonist-mediated activation mode for GPR116 in alveolar type 2 epithelial cells and alveolar homoeostasis. This is, to our knowledge, the first *in vivo* demonstration of the key role of the GPS cleavage and the unmasked tethered agonist in a mammalian system.

We set out to investigate the molecular mechanisms for GPR116 activation by its tethered peptide. The tethered agonist sequence of each aGPCR family member is highly conserved between species, which explains cross-activation among orthologues (Liebscher et al., 2014; Brown et al., 2017). In this study, we identified F995, L998, and M999 as key amino acids in the tethered agonist sequence required for GPR116 activation. These three amino acids are highly conserved in the ECDs of multiple aGPCR CTFs (Suppl Fig11), suggesting they play a critical structural role for activation in the aGPCR family. In this sense, it may explain some of the promiscous activities observed within but also between aGPCR subfamilies (Demberg et al., 2017).

Several studies analyzing the role of the amino acids composing the tethered agonist are in accordance with our identification of phenylalanine at position 3, as the most N-terminal residue playing a key role in aGPCR activation (GPR64/ADGRG2, Azimzadeh et al., 2019 and Sun et al., 2020; GPR126, Liebscher et al., 2014; Lphn3/ADGRL3, Mathiasen et al., 2020). Similarly, amino acids at positions 6 and 7, albeit not strictly conserved as leucine and methionine residues in all aGPCR tethered ligands, Leucine 998 and Methionine 999 are key for GPR116 activation. Still, it is noteworthy that additional amino acids have been have been identified to be critical for other aGPCRs possibly explaining the specificity for activation exhibited by some receptors. These studies also showed that alanine substitution of the first amino acid within the tethered agonist (usually a threonine) can lead to a loss of activity (Liebscher et al., 2014) while replacement with a hydrophobic residue, can increase the activity of the peptide (Sun et al., 2020). This is in line with our observations with the GAP10-Pro1 peptide, further supporting that this residue at position 1 can modulate the strength of molecular interactions between ligand and its binding during GPR116 activation.

With respect to the 7TM, our mutational analysis identified ECL2 as the structural component most involved in receptor activation by the tethered agonist, and we present data suggesting that ECL2 amino acids directly interact with tethered agonist residues. Specific interactions of the tethered agonist with putative target amino acids have been suggested for GPR114/ADGRG5, in which a splice variant exhibiting a single amino acid deletion in the tethered agonist does not allow for these interactions and strongly impacts receptor activation (Wilde et al., 2016). In the current work on GPR116, we identified Y1158, R1160, W1165 and L1166 as key amino acids in ECL2 and T1240 at the top of TM6 / ECL3. In particular, we noted W1165 and L1166 immediately following the cysteine involved in the disulfide bridge linking the top of TM3 and ECL2, comprising a CWL triad in which tryptophan is globally conserved in aGPCRs and the leucine (or sometimes isoleucine) is commonly present at this position (Suppl Fig11). Residues of ECL2 have also been implicated in ligand binding and activation of classical GPCRs (Wheatley et al., 2012). In particular, the CW motif in the ECL2 is conserved in family B, suggesting commonalities in the activation mechanism. In particular, the (C)WF in ECL2 is involved in CRF1 binding to CRF1R (Gkountelias et al., 2009). In PAR1 a (C)HD sequence is important for receptor activation by the peptide, further highlighting the relevance of the ECL2 amino acids following the disulfide bridge, not only for activation per se but also ligand specificity (O’Brien et al., 2001). Within the aGPCR family, Sun et al. (2020) also identified in GPR64/ADGRG2 the (C)WI triad in ECL2 and residues in TM6 as being involved in receptor activation by the tethered agonistic peptide.

The tyrosine at position 1158 is located 6 amino acids upstream of the CWL cluster and it is conserved in most aGPCRs, except in the ADGRA and ADGRG subfamily members, and ADGRV1 (Suppl Fig11). It is noteworthy that the CWL cluster is also less strictly conserved in these ADGRs. A tyrosine at a position further upstream in the ECL2 may functionally replace Y1158 in these receptors, depending on the flexibility of the ECL. R1160 is present in all ADGRFs (and GPR116 orthologs), but not conserved in other subfamilies. Hence, CWL may be involved in more fundamental and structural aspects aGPCR activation, while the tyrosine equivalent to Y1158 in GPR116 may distinguish different patterns of activation among ADGRs. R1160 in ADGRFs may be a key amino acid driving the specificity of activation by their tethered agonist sequence.

Investigating how the tethered agonist may activate GPR116 via ECL2, we identified potential amino acid pairs important for receptor activation, notably F995 / Y1158 and D1002 / R1160. The relevance of D1002 is clearly supported by the observation that a 10-amino acid peptide is much more active than GAP9, containing only the first 9 residues and lacking D1002 (Brown et al., 2017). To further validate the F995 / Y1158 interaction other approaches such as using peptides in which F995 is replaced by artificial amino acids with reduced electrons (such as Cl or F variants) may provide supporting data.

A role for ECL3 in aGPCR activation has already been suggested in the analysis of disease-associated mutations of GPR56/ADGRG1 (Kishore et al., 2017; Luo et al., 2014). The mutation L640R (corresponding to H1250 in GPR116) in ECL3 affects GPR56 signaling and specifically collagen 3-mediated signaling, probably by inducing a conformation that locks the receptor in an inactive state. In GPR116, T1240 in TM6/ECL3 may indirectly be involved in the activation mechanism. It is indeed known that the extracellular facing portions of TMs 3-6-7 come together during activation of family A GPCRs (Wheatley et al., 2012). Hence, T1240 (TM6, required for activation), as well as A1254 and V1258 (TM7, modulating potency of GAP activation), may be involved in a similar activation mechanism, transmitting the signal induced via ECL2 interactions.

During the finalization of this manuscript, a cryo-EM structure of the aGPCR GPR97/ADGRG3 was published, showing that the ECL2 contains an extended beta sheet and bends over ECL3 (Ping et al., 2021) - which is in accordance with our findings on the role of TM6- TM7 residues participating in receptor activation. The stretch of hydrophilic residues found in the ECL2 of GPR97/ADGRG3, upstream of the CWL triad, is not conserved and may provide elements of specificity distinguishing aGPCRs.

The model of the GPR116 activation that we present here is in line with other studies of aGPCRs and highlights tethered agonist:ECL2 interactions; however, it does have some limitations. The potential role of the NTF, how it may be positioned compared to the CTF and how it may affect its structure is not known so far. Modeling the highly flexible loops remains challenging, although some studies already focused on modeling of GPCR ECLs (Wink et al., 2019). In particular, ECL2 is the largest ECL in GPR116, as in many other GPCRs, and the disulfide bridge with TM3 does not introduce sufficient constraints to develop a better defined model. Even with adequate representation of the amino acid stereochemistry significant degrees of freedom remain for the ECL conformation. Of note, the cryo-EM structure of GPR97/ADGRG3 reveals a conformation of the receptor in an activation state induced by an agonist different from the tethered peptide. It will be interesting to identify the position of ECL2 in the peptide-activated state. As an extreme example, in GPR52, a class A receptor, it was recently shown that ECL2 can act as a tethered ligand and activates the receptor by interacting with the TM barrel (Lin et al., 2020). This is yet another potential mechanism of aGPCR activation and cannot be excluded for other members of the aGPCR family. Additional molecular studies, including cross-linking approaches will facilitate the identification of additional intramolecular interactions.

Our current working model postulates that residues in the tethered agonist sequence interact with amino acids located in ECL2 to induce a conformational changes resulting in productive Gq engagement and activation. While the endogenous ligand or the biological process(es) leading to displacement of the non-covalently linked N-terminal domain and subsequent GPR116 activation *in vivo* remains unknown, our data advance the mechanistic understanding of GPR116 activation in the context of pulmonary surfactant homeostasis and may facilitate the development of novel receptor modulators that can be used to treat clinically relevant lung diseases.

## Material and Methods

### 1. H991A knock-in mice

The H991A mutation was introduced into the mouse *Adgrf5* locus via CRISPR/Cas9 editing. sgRNAs with the sequence caccGGTGGAGAATGACGTCAGG were co-injected with the donor sequence listed below and CAS9 protein into fertilized C57Bl6 eggs at the 4 cell stage. Injected eggs were implanted into pseudopregnant females and tail DNA from offspring were analyzed by PCR and sequencing. A total of 9 founder pups were born from a single injection from which tail DNA was analyzed by PCR and sequencing. Three out of nine pups (lines 2552, 2553, 2556) contained the desired H991A mutation on at least one allele and either partial KI, indel or WT sequence on the other allele. These three F0 founders were bred with WT C56Bl6 mice and F1 offspring were sequenced to identify mice that carried the H991A allele. F1 mice were then bred establish stable transgenic lines for subsequent analysis. sgRNA sequence: caccGGATGGAGAATGACGTCAGG. Donor sequence (bold sequence is Ala codon to replace H991 and introduction of SpeI site, silent mutation, for screening purposes is italicized): CAATACAGGGGGCTGGGACAGCAGTGGGTGCTCTGTGGAAGATGATGGTAGGGACAATA GGGACAGAGTCTTCTGCAAGTGTAAC**gca**CTGAC*tagt*TTCTCCATCCTCATGTCCCCAGACT C

Primers for PCR screening: WT allele: FWD– CCACCTGACGTCATTCTCCA, REV-GGCGCATATAGGAAGTTCGG (product=188bp); H991A allele: FWD-CACACAGGCTGTTTCGTTGA, REV-CGCACTGACTAGTTTCTCCATC (product=298bp).

### 2. Constructs

Human and mouse GPR116 full-length constructs (hFL and mFL, respectively) have been described earlier, as well as the mouse CTF constructs (mCTF) with N-terminal mCherry or FLAG (Brown et al., 2017). The mCTF mCherry and mCTF (FLAG) constructs exhibit basal activity; the mCTFbax, derived from the latter construct by alanine mutation of the 3 N-terminal amino acids of the tethered agonist sequence, has no basal activity (Brown et al., 2017). Mutants of these constructs have been generated by site directed mutagenesis using PCR-based techniques.

A human GPR116 CTF construct deleted for the 6 N-terminal amino acids of the tethered agonist sequence and V5-tagged at its C-terminal end was generated by gene synthesis (including codon optimization; Genewiz, Leipzig, Germany) and cloning into pcDNA3.1; this construct is referred to as hCTFbax and exhibits no basal activity. The hCTFbax mBS chimera was derived from this construct, by gene synthesis. Other mutants of the human GPR116 CTF or full-length constructs were generated by site directed mutagenesis using PCR-based techniques.

The H991A-V5 cDNA construct was generated by gene synthesis (Genewiz, South Plainfield, NJ), sequenced verified and cloned into pcDNA3.1+ for expression in cultured cells. Amino acid numbering of CTF constructs is performed according to the mGPR116 sequence (Uniprot G5E8Q8). For hGPR116 CTF constructs, the mGPR116 numbering was utilized, as the CTF length is conserved in the two species, leading to a 1 to 1 alignment of amino acids.

### 3. Cells, transfection and culture

HEK293, HEK293H and HEK293T cells were obtained from ATCC (Manassas, Virginia) and used indifferently. HEK293 and HEK293H were grown in DMEM/HAMs F12 medium supplemented with 10% FCS, and 1% penicillin/streptomycin; HEK293T were grown in high glucose DMEM supplemented with 10% FCS, and 1% penicillin/streptomycin. For selection and maintenance of stably transfected cells, G418 (400 µg/ml) was added to the culture medium. Absence of mycoplasma contamination was verified.

Expression constructs were transiently transfected in HEK293 cells using Metafectene-Pro (Biontex T040-0.2; 2×10^6^ cells/well in 6-well plates, transfected with 2 µg plasmid), Lipofectamine 2000 (Invitrogen), PEImax (Polysciences) or the Amaxa Cell Line Nucleofector Kit V (Lonza VCA- 1003; 1×10^6^ cells/well in 6-well plates, transfected with 3-4 µg plasmid) according to the manufacturers protocols. Cells were seeded in the assay plate the next day and the assays were ran one day later.

For some constructs, stable populations were generated by treating the cells with G418. After complete selection of resistant cells, these were used in assays.

Stable HEK293 clones for full-length hGPR116 (clone A6) and mGPR116 (clone 3C) were described earlier (Brown et al., 2017).

### 4. Reagents

All GAP peptides were designed with a norleucine (Nle) replacing the methionine in position 7 of the tethered agonist sequence of GPR116 (e.g. GAP14: TSFSILMSPDSPDP or TSFSILNleSPDSPDP). The Nle modification was shown in Fig3E to have no major impact on the peptide activity and ensures better stability over time. Peptides were ordered at Peptides and Elephants (Hennigsdorf, Germany) or from ThermoFisher (Waltham, MA, USA) as TFA salts and dissolved as 50 mM stock solution in DMSO or in ddH_2_O for GAP10 and GAP16 experiments in Figs 1, 2 and 3C.

### 5. Expression levels of the constructs

#### Immunocytochemistry

Expression of the various GPR116 constructs tagged with V5 or FLAG was verified by immunocytochemistry (Cytofix/Cytoperm, BD 51-2091KZ). Cells were plated in 8-chamber slides (Ibidi, Cat #161107), washed the following day with PBS, fixed and permeabilized for 20 min at 4 °C, washed twice, blocked for 20 min at room temperature and stained overnight at 4 °C using an anti-V5 antibody (1/500; Invitrogen, Cat R96025) or anti-FLAG antibody (1:100; Sigma, F1365) followed by an anti-mouse secondary antibody (V5: 1 hr, RT, 1/500; Invitrogen, Cat #A21200; FLAG: 2 hr, RT, 1:200; Invitrogen, #A11004) according to the manufacturer instructions. Nuclei were stained for 3 min with Hoechst (Invitrogen, Cat #H-3570). Cells were then analyzed by confocal microscopy on a Zeiss LSM700 inverted microscope.

#### Western Blot

Transfected cells (approximately 3×10^6^ cells) were plated in a 10 cm or 6-well dish and expression was analyzed 24-48 hr after transfection, in parallel to functional assays. Cells were washed with PBS and lysed in RIPA buffer (Amresco N653-100) complemented with a protease inhibitor cocktail (Calbiochem #539137). After sonication (40%, cycle 2x for 20 seconds), total protein concentration of the cell lysates was determined by BCA assay (Pierce) and equal amounts of protein were separated by SDS-PAGE using NuPAGE® 4-12% Bis-Tris or Tris-Glycine gels. Proteins were electrophoretically transferred to PVDF membranes (Biorad #1704156) or Amersham Protran nitrocellulose membranes (Millipore, #GE10600010) and probed with a monoclonal antibody directed against V5 (clone V5-10, Sigma V8012) or FLAG (clone M2, Sigma F1365), and beta tubulin (Cell Signaling, #2128 (9F3) or actin (Seven Hills Bioreagents, clone C4, Cincinnati, OH)). Blots were then probed with donkey anti-mouse / anti-rabbit IgG HRP (Jackson ImmunoResearch 715-035-150 / 711-035-152) and analyzed for chemiluminescence on a Biorad reader.

#### Flow cytometry

The expression of mCTF mCherry constructs was quantified by flow cytometry, making use of the N-terminal red fluorescent mCherry (excitation 561 nm, emission 620 nm). Transfected cells (1×10^6^ cells) were analyzed 48 hr after transfection, in parallel with functional assays.

### 6. Calcium transient assays

HEK293 cells were seeded in 384-well plates (Greiner, #781946) at 30 000 cells/well. Medium was removed the following day and cells were loaded for 2 hr at 37 °C, 5 % CO2 with 20 µl Calcium 6 loading buffer (Molecular Devices, Cat #R8191): 1x HBSS (Gibco, Cat #14065.049) containing 20 mM HEPES (Gibco, Cat #15630) and 2.5 mM probenecid (Sigma, Cat #P8761). Plates were then transferred to a FDSS7000 fluorescence plate reader (HAMAMATSU, Japan) and Ca^2+^ responses were measured in real time using a CCD camera with excitation 480 nm/emission 540 nm. Prior to stimulation, baseline measurements were recorded over 5 sec (Fmin). For stimulation, 20 µl of test compounds prepared 2x in HBSS containing 20 mM HEPES were dispensed. Fluorescence was read over 1-1.5 min. Maximum fluorescence (Fmax) was determined and normalized to baseline. Data are reported as ΔF/F = (Fmax – Fmin)/Fmin. Adequate response of the endogenous M3R was verified for each transfection.

### 7. Calcium imaging of mouse H991A alveolar type 2 cells

Primary mouse AT2 cells were isolated from homozygous H991A adult (mice 6-8 weeks) and cultured in BEGM plus 10% charcoal-stripped FBS and recombinant human KGF (10 ng/ml, Peprotech) on top of a thin substrate of 70% Cultrex (Trevigen)/30% rat tail collagen (Advanced Biomatrix) coated onto 35-mM dishes with a coverslip bottom (MatTek) to maintain a differentiated phenotype. Mouse cells were cultured for 6 days after isolation prior to experimentation. On the day of experimentation, media were replaced with 1 ml Krebs-Ringer’s buffer containing 8 mM calcium chloride and incubated for 30 min at 37 °C to equilibrate. Cells were washed once with Krebs-Ringer’s buffer containing 1.8 mM calcium chloride and loaded with Fluo-4 AM (1 μg/μl; ThermoFisher) plus 10 μl 0.2 % pluronic acid for 30 min at 37 °C. Cells were washed 3 times with 1 ml with Krebs-Ringer’s plus calcium buffer, leaving the last 1 ml on cells to image. Baseline fluorescent data were recorded for 5 min prior to addition of SCR10 or GAP10 peptides. Ionomycin (10 µM) was added 2 min prior to end of imaging as positive control. Time-lapse imaging was performed on Zeiss 200M wide-field fluorescent microscope. Data were analyzed using Slidebook software.

### 8. Inositol phosphate accumulation assays

#### Radiometric method

HEK293 cells were seeded in 12-well plates at a density of 1.5×10^5^ cells per well and transiently transfected with plasmids expressing WT, WT mGPR116-FLAG, CTF-FLAG or alanine mutants of CTF-FLAG. Twenty-four hrs post transfection, media was removed and replaced with serum-free Modified Eagles Medium (MEM; Mediatech) containing 1 µCi/ml of 3H-myo-inositol (PerkinElmer Life Sciences). Forty-eight hrs post-infection, media was removed and replaced with serum free media supplemented with 20 mM LiCl for 3 hr. For samples to be stimulated with GAPs, the peptides were added at the appropriate concentration when the media was replaced with that supplemented with LiCl. Reactions were stopped by aspirating medium, adding 1 ml of 0.4 M perchloric acid, and cooling undisturbed at 4 °C for 5 min. 800 µl of supernatant was neutralized with 400 µl of 0.72 M KOH/0.6 M KHCO3 and subjected to centrifugation. 1 ml of supernatant was diluted with 3 ml of distilled H2O and applied to freshly prepared Dowex columns (AG1-X8; Bio-Rad). Columns were washed twice with distilled H2O, total inositol phosphates (IP) were eluted with 4.0 ml of 0.1 M formic acid and 1 M ammonium formate, and eluates containing accumulated inositol phosphates were counted in a liquid scintillation counter. 50 μl of neutralized supernatant was counted in a liquid scintillation counter to measure total incorporated 3H-myo-inositol. Data are expressed as accumulated inositol phosphate over total incorporated 3H-myo-inositol. Adequate response of the endogenous M3R was verified for each transfection.

#### HTRF kit

IP1 accumulation was measured using the manufacturer’s instructions (IP-One, Cisbio 62IPAPEC). Briefly, cells were seeded at 60 000 cells per well in 384-well plate (Greiner #781080) one day before the assay. Plates were flicked and cells were incubated with 20 μl peptide diluted in HBSS buffer (Invitrogen 14065-049) complemented with 20 mM HEPES (Invitrogen 15630-056) and 50 mM LiCl. After 2 hr incubation at 37°C, 5% CO2, 5 μl IP-d2 in lysis buffer and 5 μl IP-Cryptate in lysis buffer were added for 1 hr at RT. Plates were read at 665 and 620 nm on an EnVision® Xcite Multilabel reader (PerkinElmer, # 2104-0020). For data analysis, the 665 nm/620 nm ratio was used as data point value and IP1 formation was measured according to the calibration curve.

### 9. Histology of H991A mouse lungs

Adult mouse (4.5 months of age) lungs were inflation fixed with 4% PFA at 20cm and fixed overnight at 4C. Tissues were dehydrated the following day and embedded in paraffin. Sections of 7 um were cut, stained with H&E and imaged on a X scope.

### 10. Snake plots

Snakeplots were generated using the web application Protter - visualize proteoforms (Omasits et al., 2013). TM boundaries are as defined in the model.

### 11. Modeling

Positions 991-1270 of the UniProt entry Q8IFZF2 (human GPR116) served as query for a Blast sequence search of the Protein Data Bank (PDB). This search resulted in two hits: PDB entries 4K5Y (Corticotropin-Releasing Factor 1 receptor (CRF1R) in complex with an antagonist, 27.9% sequence identity) and 4Z9G (CRF1R in complex with an antagonist, 28.5% sequence identity). The CRF1R receptor belongs to the class B1 (secretin) of the GPCR superfamily. Only class B1 shares some sequence similarity with aGPCRs (de Graaf et al., 2016). We selected the PDB entry 4K5Y which exhibits a slightly better resolution than 4Z9G, as template for homology modelling of the 7TM domain of human GPR116.

We used Maestro 2019-1 (Schrödinger) for all the molecular modelling tasks. To start with, the lysozyme T4 insert was removed from the PDB entry 4K5Y. This curated PDB entry served as reference for a pairwise sequence alignment of the 991-1270 of the UniProt entry Q8IFZF2, using the Schrödinger implementation of the BLOSUM62 matrix and default settings. Because of the low sequence identity between CRF1R and the 7TM domain of GPR116, we applied several manual corrections to the resulting alignments to remove gaps in secondary structure elements and improve the alignment of conserved motifs in the 7TM domains of adhesion and secretin family GPCRs. The corrected alignment served as input for the homology modeling module of Prime, using default settings for both the structurally conserved regions and the ECLs. We modelled all the ECLs but ICL2, which is missing in the template. The resulting model was subsequently submitted to the Protein Reliability Report module of Maestro 2019-1. Its quality was deemed satisfactory.

We used the Select Rotamers module of Maestro 2019-1 to explore the different side chain conformations of selected amino acids and Schrödinger’s SiteMap module was run to identify putative binding sites in the homology model (SiteMap, version 2.4, Schrödinger, LLC, New York, NY, 2010).

## Acknowledgements

We would like to thank the CCHMC gene editing core for their efficient support and Johannes Voshol (Novartis) for helpful discussions throughout this work.

## Supplementary Figures

**Supplemental Figure 1:**
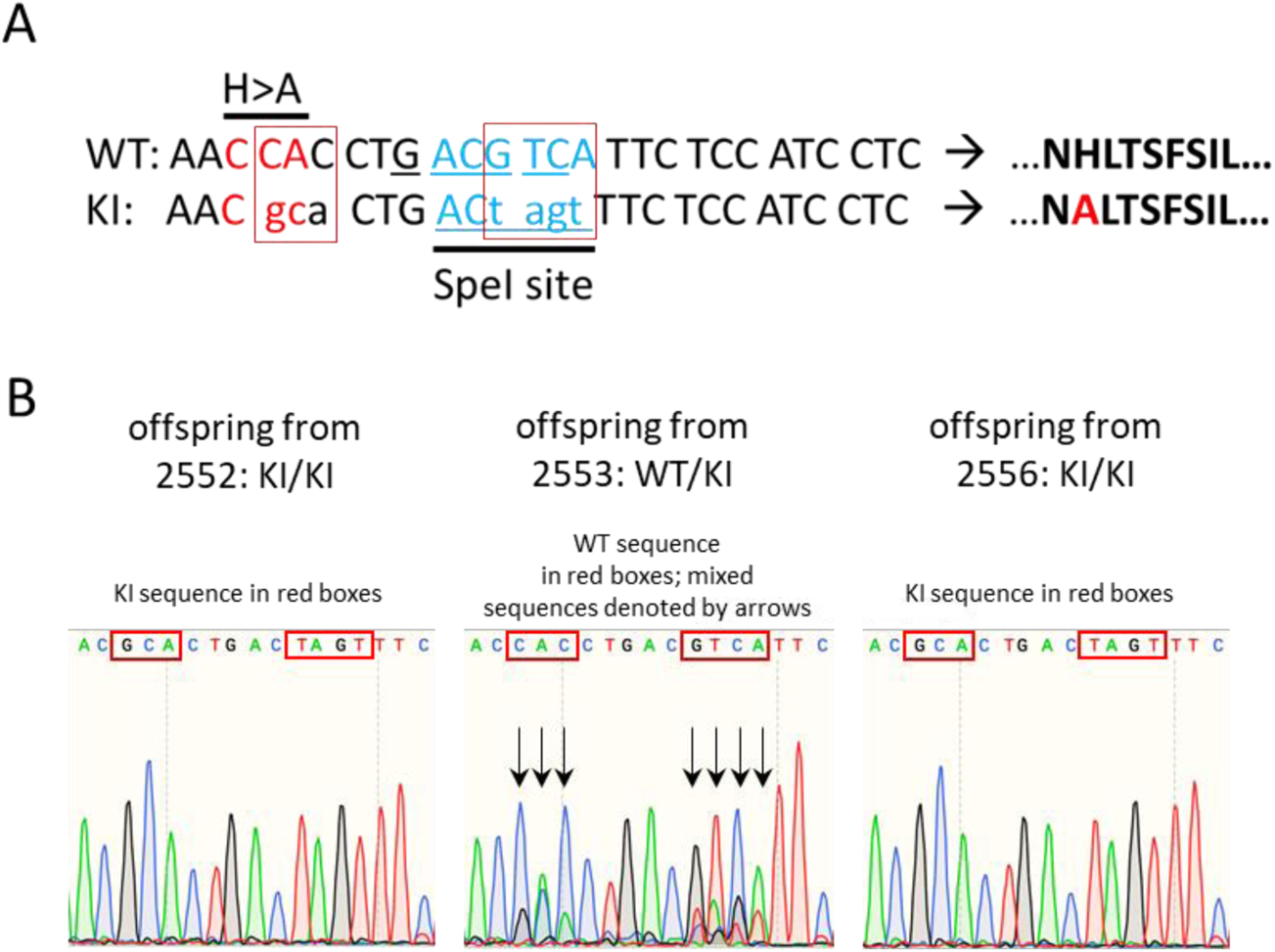
Schematic of H991A point mutation introduced into the Adgrf5 locus via CRISPR/Cas9 gene editing. **A.** Red boxes denote introduced base pair changes in the Adgrf5 locus. SpeI restriction enzyme recognition site was introduced for screening purposes without altering amino acid sequence at this position. **B**. Representative sequencing traces of genomic DNA from two homozygous (2552 and 2556) and one heterozygous H991A line (2553). Line 2553 was bred to homozygosity for experiments.

**Supplemental Figure 2:**
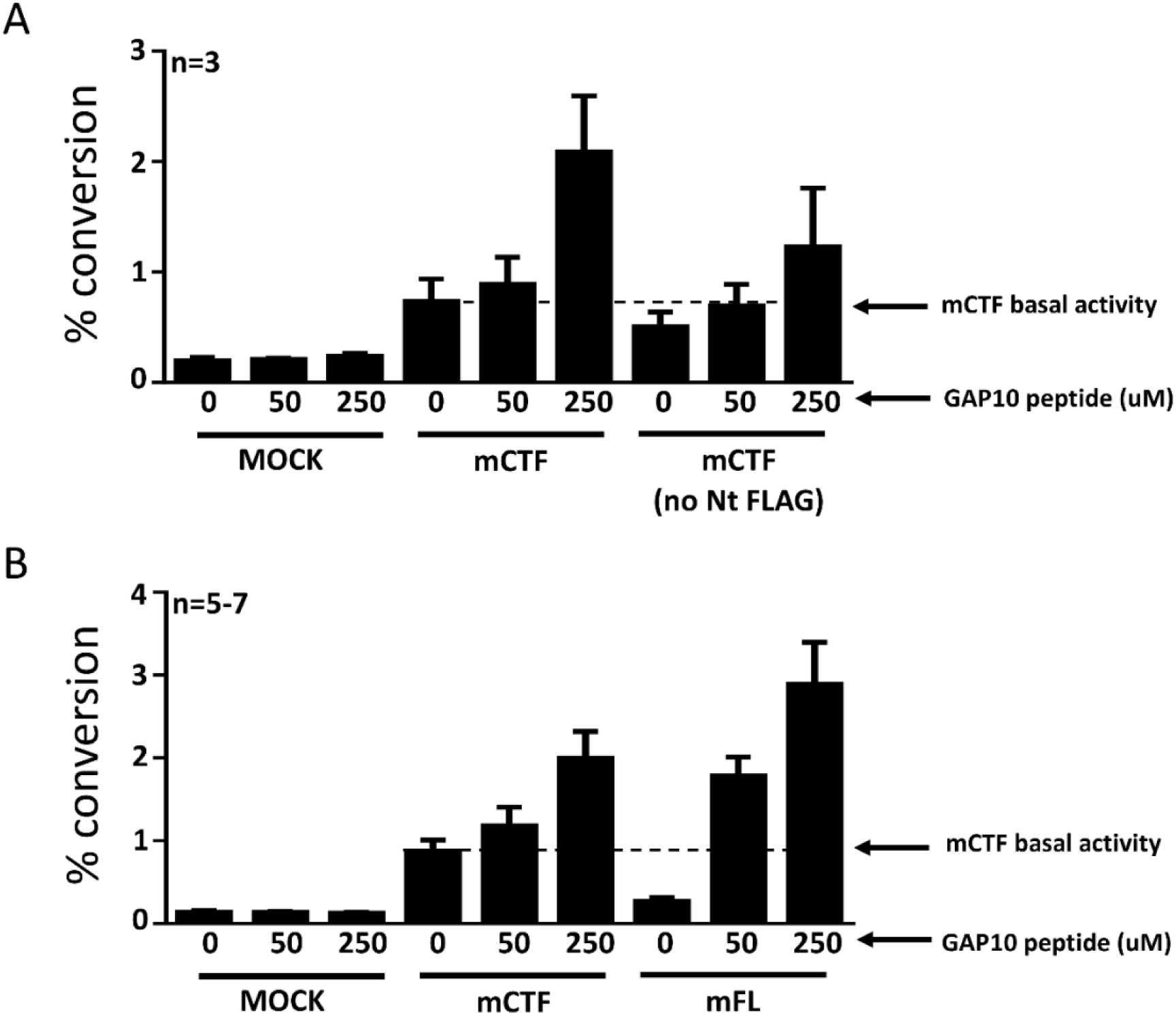
Super-activation of mGPR116 CTF constructs with basal activity. **A.** The N-terminal FLAG tag in the mCTF constructs does not affect the activation by GAP10. Tagged and untagged mCTF were transiently expressed in HEK293 cells and GAP10-induced IP1 conversion was measured. Dotted line indicates mCTF basal activity; n=3. **B.** The mCTF construct, expressed transiently in HEK293 cells, shows basal activity and can be super-activated to levels approximating that of the full-length receptor (mFL) upon addition of GAP10. Receptor signaling is measured in the IP1 conversion assay; n=5-7.

**Supplemental Figure 3.**
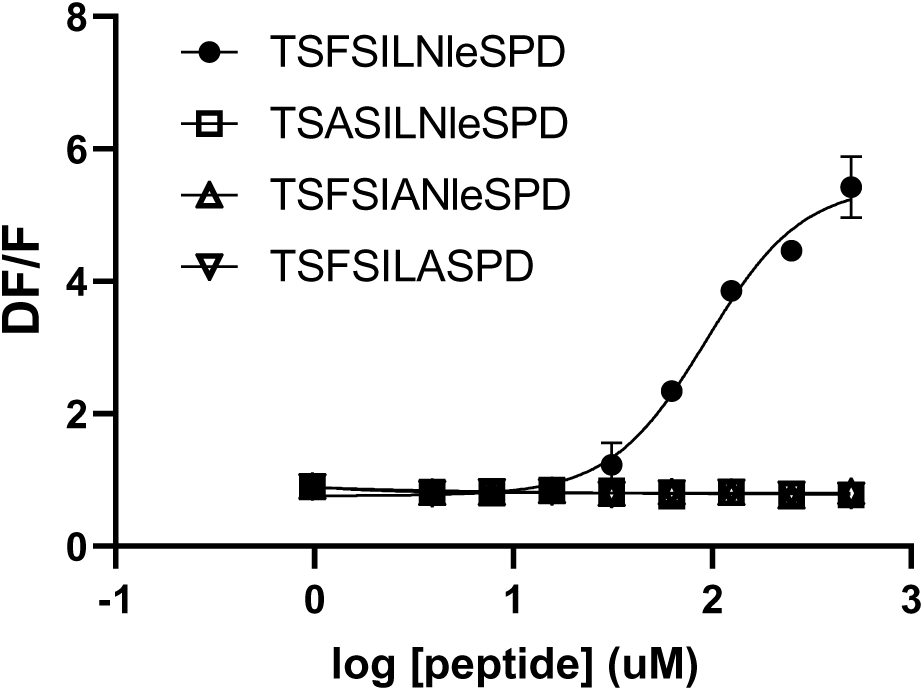
Alanine mutation of key amino acids in GAP10 inactivates the peptide for human GPR116 full-length stimulation. Full-length hGPR116 (HEK293 clone A6) stimulated with GAP10 and peptide variants corresponding to alanine changes identified in mGPR116 as being critical for the activation by the tethered agonist. (n=3 independent experiments, 4 biological replicates per group).

**Supplemental Figure 4.**
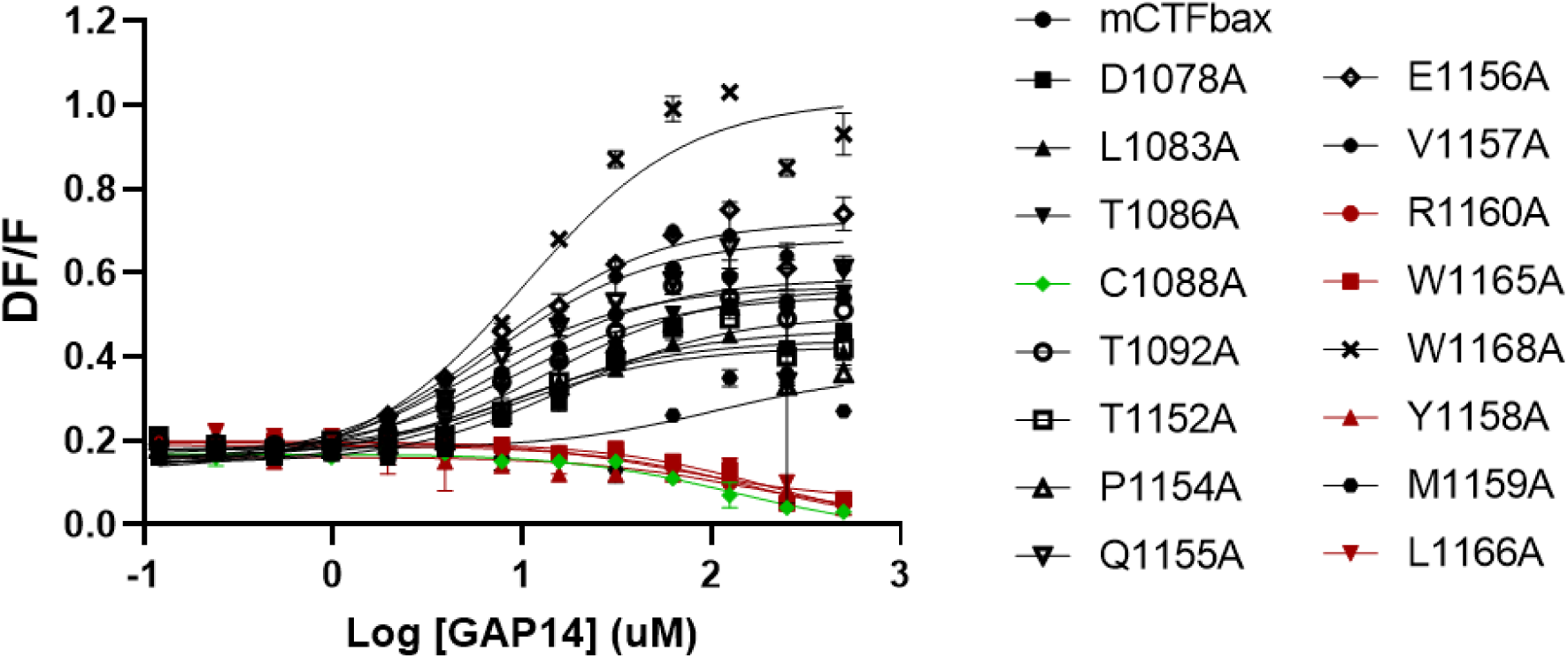
Alanine scan of GPR116 CTF ECLs – representative data from mutants with or without functional effect. Calcium transient assays were performed in transiently transfected HEK293 cells with exogenous GAP14 as the stimulus. mCTFbax mutants with varying effects of the mutation are shown. Mutants exhibiting no activity upon GAP14 stimulation and selected for further analysis are highlighted in red and the construct with one of the cysteines from the disulfide bridge mutated to alanine is in green. Data are from one experiment with biological 4 replicates are shown.

**Supplemental Figure 5.**
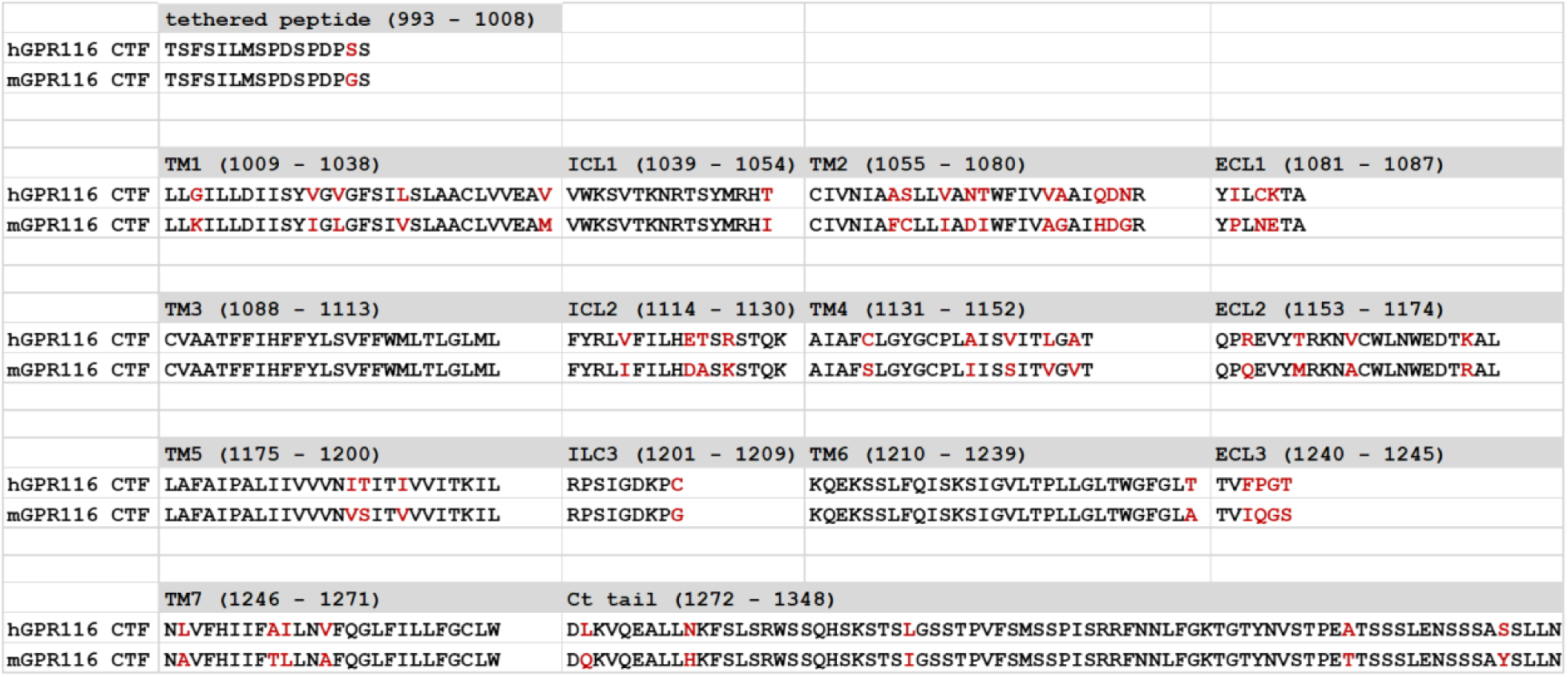
Alignment of the human and mouse GPR116 CTF sequences. Residues that are not conserved between the 2 species are highlighted in red.

**Supplemental Figure 6.**
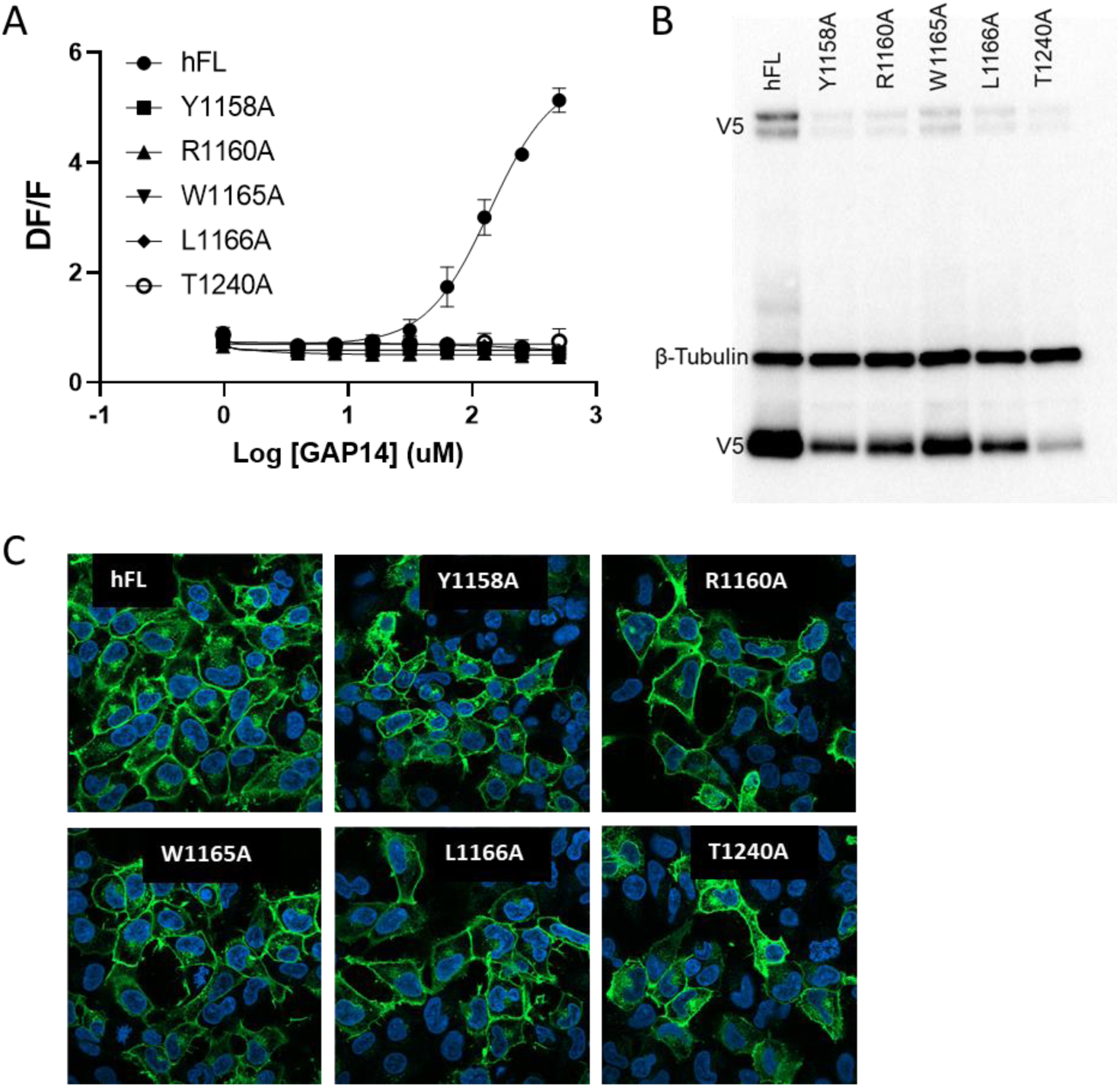
Mutation of the key ECL amino acids in human GPR116 full-length leads to inactive constructs. **A.** Activity of the full-length hGPR116 mutated for key ECL2/TM6 residues, as compared to the WT hGPR116 A6 clone (hFL) . Mutants were stably expressed in HEK293 cells and GAP14-induced calcium transients were measured. Representative data of 3 repeat experiments are shown. **B.** Expression of hFL WT (clone A6) and mutants in stably transfected HEK293 cells. Constructs were detected by Western blot using an anti-V5 antibody. The major band (∼30kD) corresponds to the cleaved CTF and the minor bands at high molecular weight (∼185kDa) correspond to the full-length receptor. **C.** Membrane expression of hFL WT (clone A6) and mutants. The expression of the mutants as HEK296 stable populations was tested by V5 tag immunocytochemistry.

**Supplemental Figure 7:**
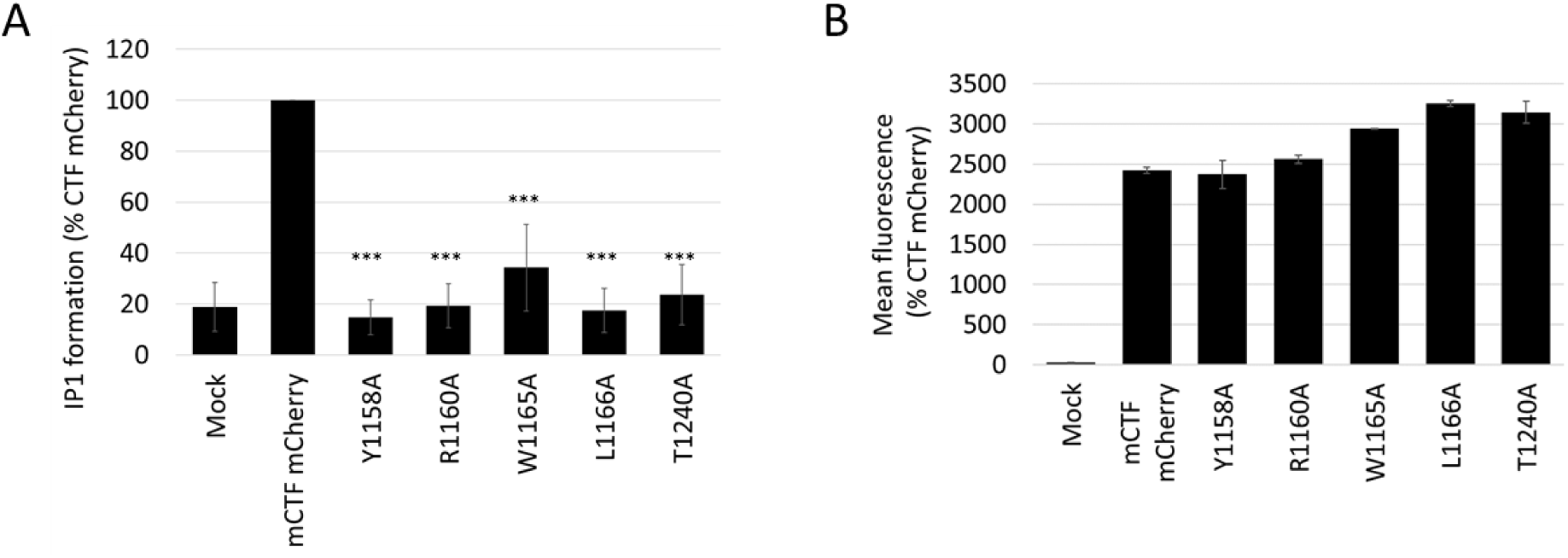
Alanine mutation of key ECL residues in mGPR116 CTF constructs with basal activity. **A.** Basal activity of ECL mutants in the mCTF mCherry parent construct. ECL residues important for activation were mutated to alanine and IP1 formation was measured in transiently transfected HEK293 cells. Data, normalized to the parent construct, are expressed as mean +/- SD (n=8 repeat experiments; 1-way ANOVA, *** P<0.0001). **B.** mCTF mCherry constructs were transiently expressed in HEK293 cells and expression was evaluated by flow cytometry analysis of the mCherry signal.

**Supplemental Figure 8.**
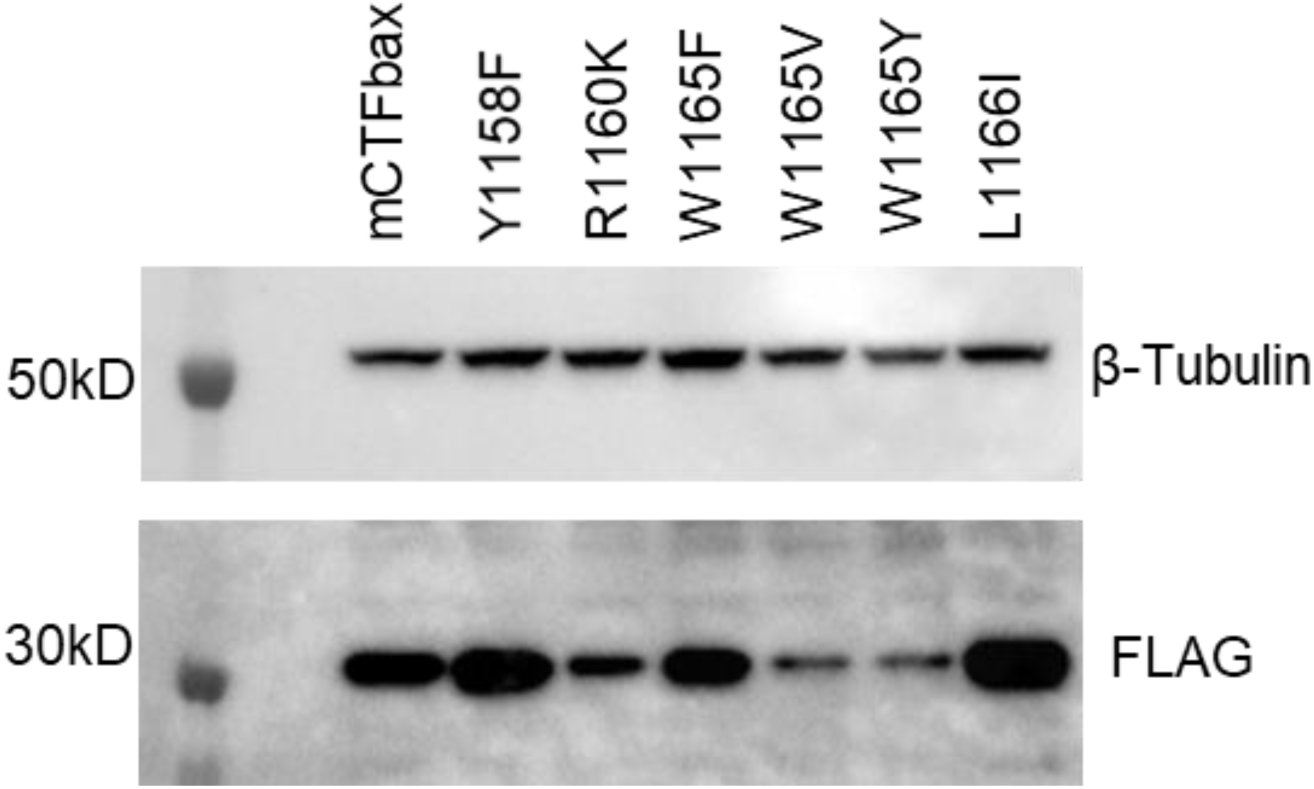
Expression of mCTFbax mutants in HEK293 stable populations. Expression levels were analysed by Western Blot using the FLAG tag

**Supplemental Figure 9.**
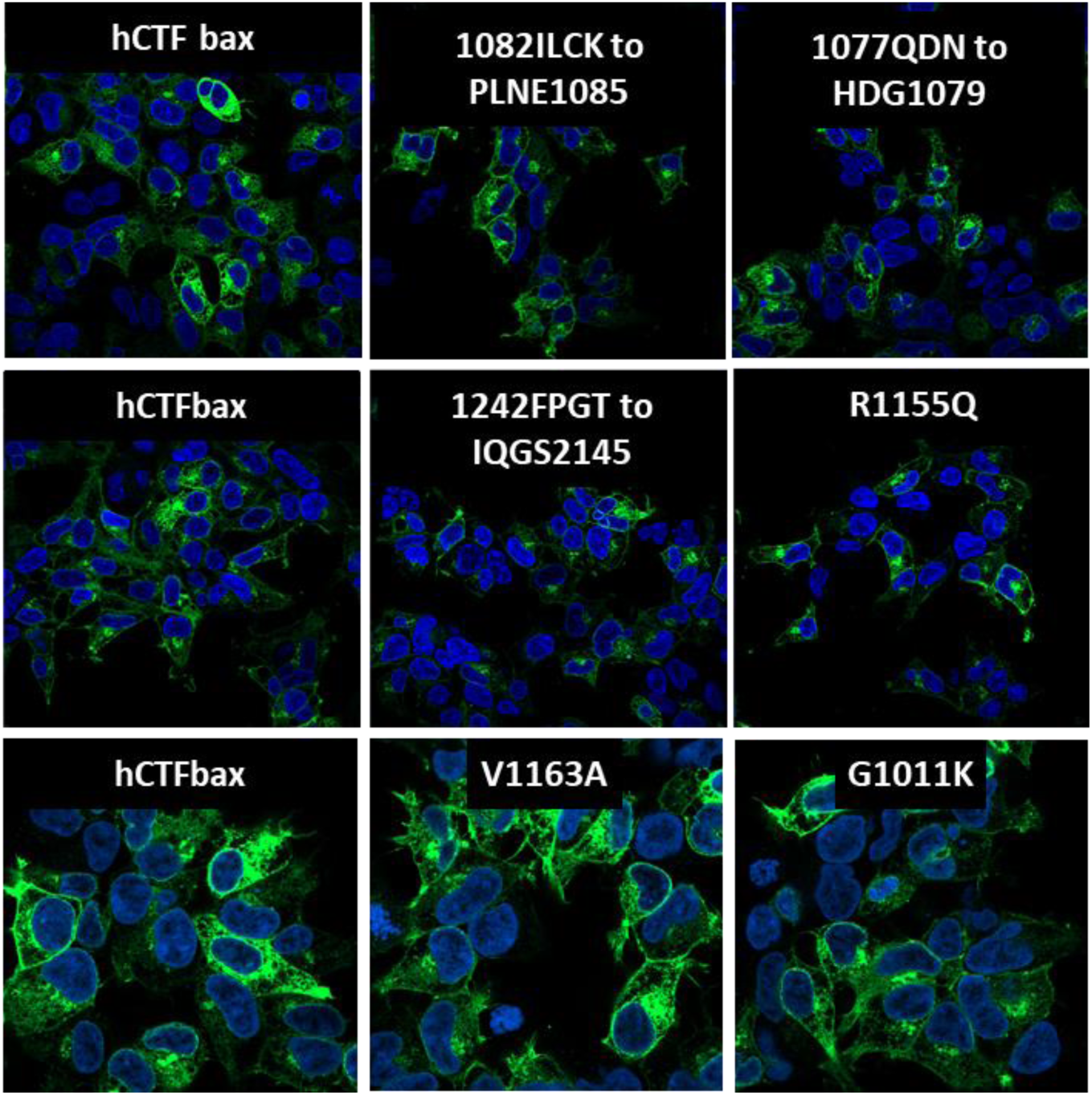
Evaluation of the role of ECL and N-terminus amino acids not conserved between human and mouse GPR116 CTF. Expression of hCTFbax mutants as HEK293 stable populations was tested by V5 tag immunocytochemistry.

**Supplemental Figure 10.**
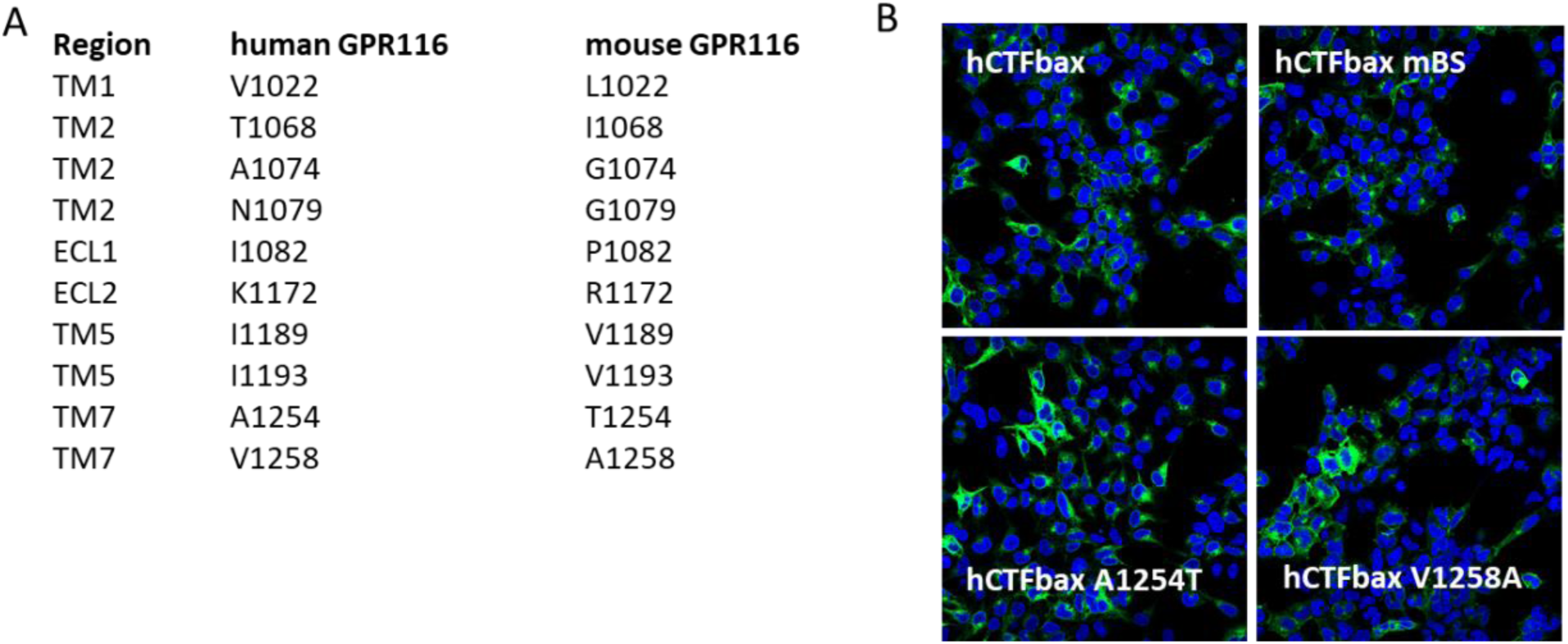
Details of the putative binding site and expression of corresponding mutants. **A.** Amino acids defining the potential binding site in mouse GPR116 (mBS). Corresponding residues in human GPR116 are given. **B.** Expression of hCTFbax, the related hCTFbax mBS, A1254T and V1258A mutants in HEK293 cells. The C-terminally tagged constructs are detected by immunocytochemistry using an anti-V5 antibody.

**Supplemental Figure 11.**
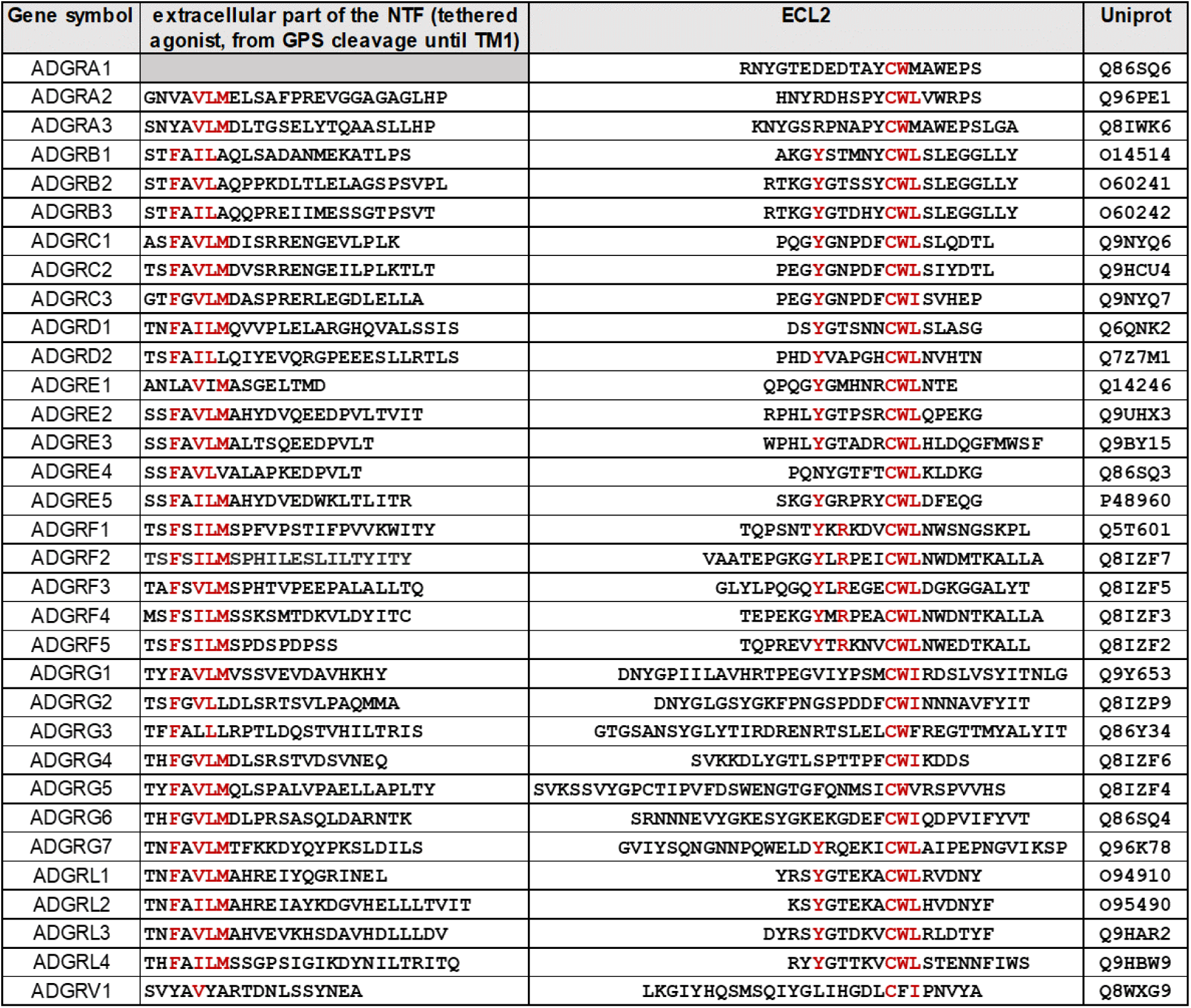
Alignment of the human aGPCR tethered agonist and ECL2 sequences. Conserved residues mentioned in the text are highlighted in red.

